# LineageVAE: Reconstructing Historical Cell States and Transcriptomes toward Unobserved Progenitors

**DOI:** 10.1101/2024.02.16.580598

**Authors:** Koichiro Majima, Yasuhiro Kojima, Kodai Minoura, Ko Abe, Haruka Hirose, Teppei Shimamura

**Affiliations:** Department of Systems Biology, Nagoya University Graduate School of Medicine; Laboratory of Computational Life Science, National Cancer Center Research Institute; Japanese Red Cross Aichi Medical Center Nagoya Daiichi Hospital; Division of Computational and Systems Biology, Tokyo Medical and Dental University Medical Research Institute

**Keywords:** single-cell RNA sequencing (scRNA-seq), lineage tracing, machine learning, deep learning, neural network, deep generative model, variational autoencoder (VAE), dimensionality reduction, time series data, differentiation, hematopoiesis, reprogramming, tumor evolution, generalized linear model (GLM), transcription factor activity

## Abstract

Single-cell RNA sequencing (scRNA-seq) enables comprehensive characterization of the cell state. However, its destructive nature prohibits measuring gene expression changes during dynamic processes such as embryogenesis. Although recent studies integrating scRNA-seq with lineage tracing have provided clonal insights between progenitor and mature cells, challenges remain. Because of their experimental nature, observations are sparse, and cells observed in the early state are not the exact progenitors of cells observed at later time points. To overcome these limitations, we developed LineageVAE, a novel computational methodology that utilizes deep learning based on the property that cells sharing barcodes have identical progenitors. This approach transforms scRNA-seq observations with an identical lineage barcode into sequential trajectories toward a common progenitor in a latent cell state space. Using hematopoiesis and reprogrammed fibroblast datasets, we demonstrate the capability of LineageVAE to reconstruct unobservable cell state transitions, historical transcriptome, and regulatory dynamics toward progenitor cell states at single-cell resolution.

## 1 Motivation

Single-cell RNA sequencing (scRNA-seq) is crucial in identifying cell types, understanding cellular diversity, and exploring changes in gene expression during differentiation and in response to stimuli. However, it has limitations in tracking cell behavior over time because it is a destructive assay. On the other hand, in research on stem cells, particularly induced pluripotent stem cells (iPS cells), and cancer, it is essential to explore the initial state of differentiated cells in order to discover therapeutic targets during carcinogenesis and realize regenerative medicine to restore lost functions and prevent diseases by gaining a deeper understanding of the precise mechanisms underlying cancer initiation from a single transformed cell. Recognizing the initial cellular states and early development is essential in both of these research areas. There is a need to understand cell state transitions, particularly from progenitors to differentiated states. Existing methods have limitations, and inferring progenitor cell states remains a challenge. Recent studies have shown promise in integrating scRNA-seq with lineage tracing to track clonal relationships and cell state transitions during differentiation. Variational autoencoder (VAE)-based methods effectively capture biological features; however, challenges remain in consistently inferring cell state transitions. In this study, we aimed to fill this gap by leveraging VAE-based deep learning to infer continuous dynamics, progenitor cell states, and cell fate. This contributes to a better understanding of the differentiation processes. Ultimately, we aimed to reveal the molecular factors governing single-cell transcriptome dynamics during differentiation, thus enhancing our understanding of the differentiation process. This process involves the restoration of progenitor and intermediate cell states, historical gene expression, and transcription factor (TF) activity along real-time evolution.

## 2 Introduction

Cells differentiate into various distinct phenotypes in continuous biological processes such as tissue development and disease progression. Differentiation studies aim to understand cell state transitions throughout this process, from the progenitor state to the post-differentiation state. This involves identifying the driving factors that regulate these changes and elucidating their underlying mechanisms. Single-cell RNA sequencing (scRNA-seq) has significantly contributed to our understanding of these biological processes by analyzing individual cell transcriptomes and revealing cellular heterogeneity and gene expression profiles within subpopulations (Hwang *et al*., 2018). However, scRNA-seq is limited to capturing transient snapshots of cell transcriptomes because of the destructive nature of the process, allowing each cell to be measured only once during the analysis. This limitation hinders monitoring of progenitor states and dynamic transitions between cellular states during differentiation.

Several computational methods have been developed to elucidate the cell state transitions and differentiation pathways over time. One method is the trajectory inference, which started with the concept of pseudotime (Trapnell *et al*., 2014) and has evolved into more than 40 different methods (Saelens *et al*., 2019). These methods enable the ordering of cell types along developmental trajectories in dynamic processes, such as the immune response and pancreatic beta cell maturation (Song *et al*., 2019). However, they primarily arranged cells based on the similarity of expression patterns from a single predefined progenitor state, thus failing to address the heterogeneity of progenitors before differentiation. Another approach for inferring cell state transitions during differentiation is through optimal transport calculations, such as the Waddington-OT (Schiebinger *et al*., 2019). These approaches establish connections between multiple time points, enabling inference of long-term transitions by combining transport maps between intermediate time points. This approach is based on proximity in either a high-dimensional expression space or compressed versions. Consistently capturing the process of stem cells with similar cell states differentiating into multiple cell types, such as hematopoiesis, makes it challenging.

Additionally, wet experimental methods, called lineage tracing, have been developed for differentiation analysis. Recent advancements have demonstrated that integrating scRNA-seq measurements with lineage tracing using cell barcodes allows the tracking of clonal relationships and observation of transitional cell states during differentiation at discrete and sparse time points (Weinreb *et al*., 2020; Oren et al., 2021). However, the earliest observable cells are not true progenitor cells owing to the destructive nature of the measurement and the necessity of multiple cell divisions to introduce DNA barcodes. Several methods such as LineageOT (Wang SW *et al*., 2022), CoSpar (Forrow *et al*., 2021), and PhyloVelo (Wang K *et al*., 2023) have linked observations across multiple time points through lineage relationships. However, no method has effectively exploited the shared progenitor traits of clones with identical barcodes from unobservable progenitor cell states.

To overcome these limitations, we introduced LineageVAE, which uses a variational autoencoder (VAE) to infer progenitor cell states and continuous differentiation trajectories at the single-cell level. This method considers cell state transitions from differentiated cells to progenitor cells in a latent cell state space generated by the VAE (Kingma and Welling., 2013), to recover differentiation dynamics from a common progenitor cell to cells with shared lineage information. LineageVAE depicts sequential cell state transitions from simple snapshots and infers cell states over time. Moreover, it generates transcriptomes at each time point using a decoder. To the best of our knowledge, LineageVAE is the first method that utilizes the property that the progenitors of cells introduced with a shared barcode are identical, allowing the reconstruction of historical cell states and their expression profiles from the observed time point toward these progenitor cells under the constraint that the cell state of each lineage converges to the progenitor state. This methodology enabled us to infer sequential cell state dynamics, elucidate unobservable progenitor heterogeneity and bias, reconstruct historical gene expression, and determine transcription factor (TF) activity along real-time evolution. Applied to scRNA-seq data with lineage tracing during hematopoiesis and direct reprogramming, LineageVAE has demonstrated its capability to restore backward cell state transitions toward progenitor cell states and regulatory activity along differentiation trajectories.

## 3 Results

### 3.1 The LineageVAE model

LineageVAE is based on a probabilistic generative model that assumes high-dimensional single-cell transcriptomes are derived from low-dimensional latent cell states. Latent cell states of each cell are derived from the sequential cell state transitions of progenitor cells which are shared by cells belonging to the same Lineage with identical barcodes. For the unobserved variables, latent cell states, and the cell state trajectory toward the corresponding progenitor cells, we conducted variational inference using deep learning techniques, similar to the VAE. In LineageVAE, we assumed the following probabilistic model. As a generative model, for each lineage *l* integrated with the same DNA barcode, we assumed a common progenitor cell state ***z***_0_. At the initial cell state transition, it branches into as many cell states as the cells observed with the corresponding barcode. Each cell *i* transitions its cell state 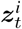 sequentially in the latent space as time passes. At time point *T*^*i*^, observational data are generated from each latent state of each cell 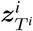. Observational data consists of scRNA-seq transcriptome measurements 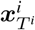 and lineage barcode *l*. Cell state transitions are modeled using the normal distribution, and transcriptome measurements 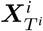 is modeled using the Poisson distribution.

For the generative model above, we repeatedly perform variational inference backward in time and estimate the transition dynamics in the latent space. For each lineage, each cell transitions its cell state from the observed time point *T*^*i*^cell state 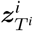 toward the common progenitor cell state ***z***_0_ sequentially in the latent space (Supplementary Fig. 1). At time point *t* = 0. the cell states of cells belonging to identical lineages converge to a common progenitor cellular state ***z***_0_.

The model was optimized as follows: First, LineageVAE uses an encoder-decoder pair to map data ***X*** to a lower-dimensional latent space ***Z***. The latent state of cell *i* at time *t* is denoted as 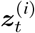, which represents a set of low-dimensional vectors of latent variables (here, set to 10 dimensions). The encoders was used to infer the variational posterior *q*_*ϕ*_(***z***|***x***) from which ***z*** is sampled, while decoders calculate the parameters of Poisson distributions, which can be written as *p*_*θ*_(***x***|***z***). Here, *ϕ* and *θ* denote the parameters for the encoder and decoder, respectively. Second, another encoder is used to estimate the dynamics, which is the difference between the latent states at a certain time and the latent state at the previous time, with *ω* denoting the parameters for the encoder for inferring dynamics. This encoder was used to infer the variational posterior:

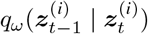

These three neural networks were optimized by maximizing the Evidence Lower Bound (*ELBO*) for each lineage:

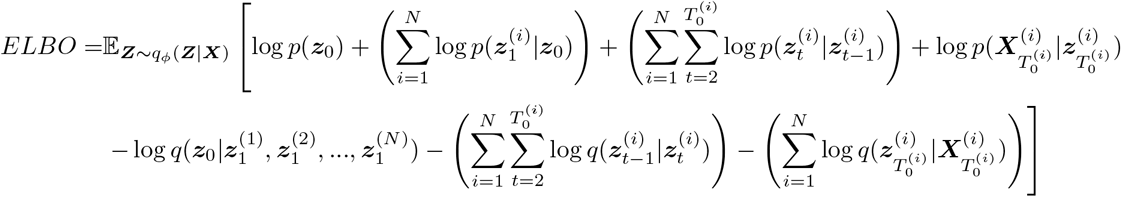

The unique training procedure of LineageVAE, which involves three neural networks, can infer unobserved progenitor cell states and historical cell state transitions between intermediate time points at the single-cell level. As an application of this method, LineageVAE allows us to elucidate progenitor heterogeneity, reconstruct historical gene expression, and estimate TF activity at each time point (Fig. 1).

**Figure 1:**
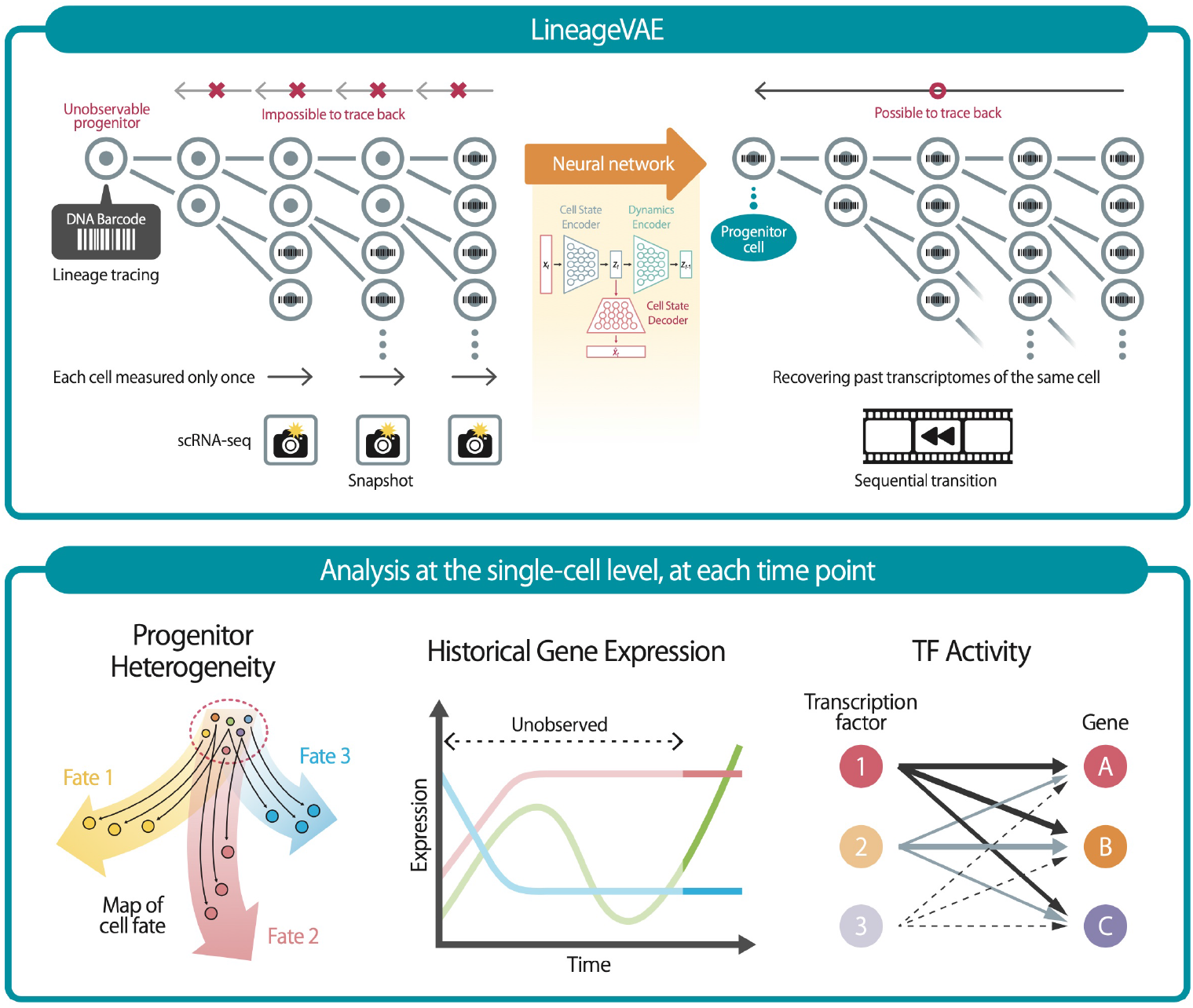
LineageVAE conducts variational inference backward in time within the latent space. **(upper)**, LineageVAE inference originates from the observation time point and extends toward the progenitor, representing the uniquely identical state of the progenitor cell within the same lineage. This enables the inference of cell states at progenitor and intermediate time points. **(lower)**, Downstream analysis enabled by LineageVAE serves various applications: i) elucidating progenitor heterogeneity, ii) reconstructing historical gene expression, and iii) estimating TF activity at each real-time point.

### 3.2 LineageVAE extracts biologically meaningful latent variables from single-cell transcriptomes with lineage tracing

To validate the performance of LineageVAE in analyzing the single-cell transcriptome with lineage tracing data, we applied LineageVAE to a recently published scRNA-seq with lineage tracing dataset of hematopoiesis, with 87,449 clones of 130,887 cells (Weinreb *et al*., 2020). The dataset included observations on day 2, 4, and 6, excluding observations on days 0, 1, 3, and 5. Despite the large number of cells in the dataset, the number of multi-cell clones and state-fate clones containing progenitors and differentiated cells, was limited. To address this limitation, we extracted lineages containing more than 20 cells and included the day 2 observational data. We set the batch size for dynamic estimation to 20, such that each lineage included as many cells as possible and as many cell types as possible, which were already annotated in the original paper (Supplementary Table 1). This is because LineageVAE learns the lineage-specific dynamics in each minibatch. We first learned the latent space using all 130,887 cells and subsequently estimated the dynamics for each selected lineage. LineageVAE embedded cells from the same annotated subpopulation with similar expression patterns closely in the latent space (Fig. 2b, Supplementary Table 2).

**Figure 2:**
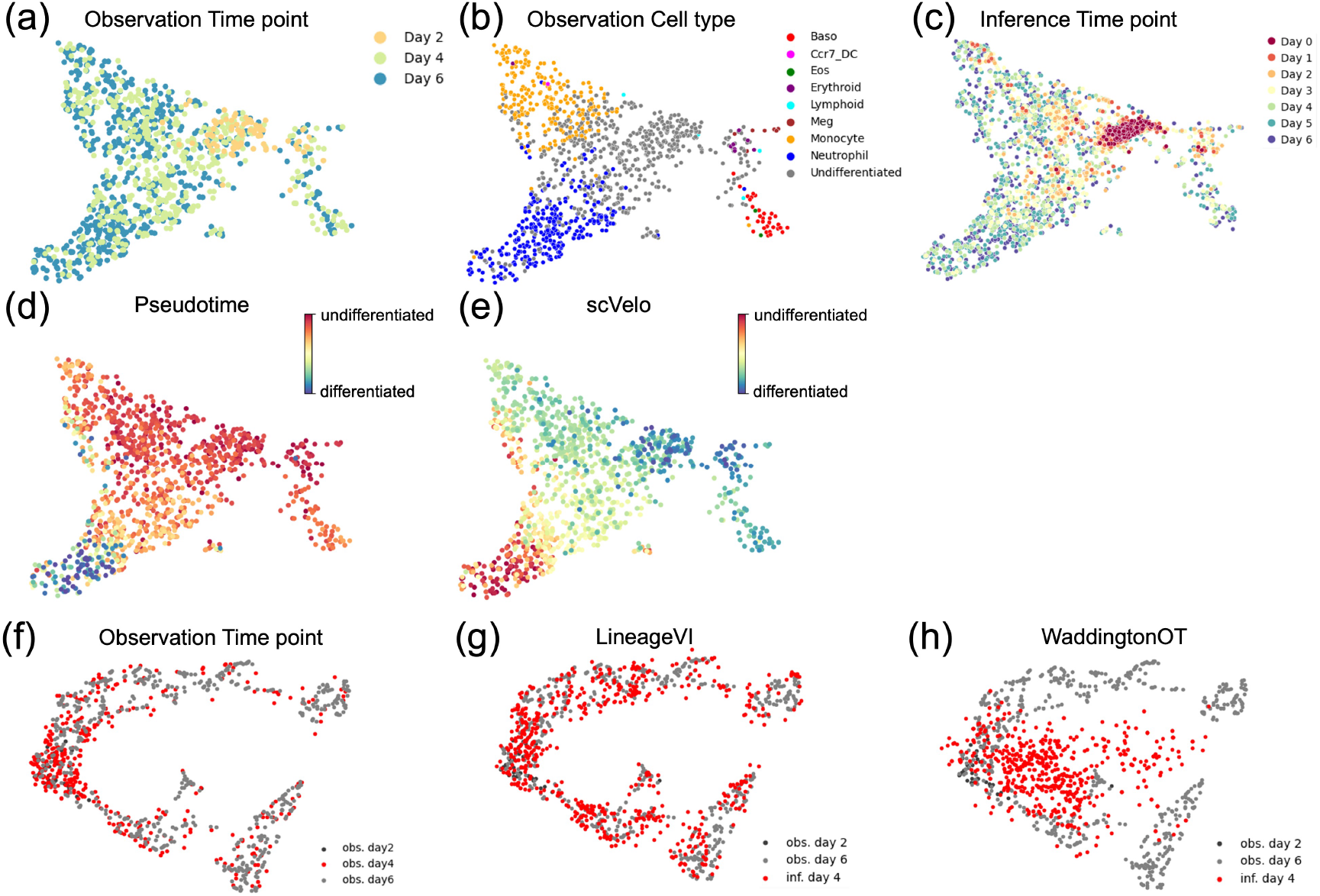
LineageVAE analysis on the hematopoiesis lineage tracing dataset. **a-c**, UMAP visualization of the latent space inferred by the model. Each point color is displayed by **a**, Observed time points. **b**, Celltype annotations. **c**, Time-series transition of cell state estimated by LineageVAE. **d**, A projection of the times inferred by the pseudotime algorithm into the same space. **e**, A projection of the times inferred by the scVelo algorithm into the same space. **f-h**, Comparison with other methods by interpolation task. **f**, observed time point. **g**, day 4 interpolation by LineageVAE. **h**, day 4 interpolation by Waddington-OT.

To assess how well the latent space reflected the biological meaning of the cells, we evaluated the accuracy of this embedding. We calculated the average coordinates and standard deviations in the latent space for each manually inspected and annotated subpopulation based on marker genes. Subsequently, we computed the Euclidean distance and p-value between the subpopulations. Our analysis confirmed that cells of the same manually annotated cell type existed in close proximity within the latent space, demonstrating that each cell type formed a distinct population (Supplementary Table 2). This shows that the LineageVAE’s latent space is biologically meaningful and captures the biological variation among cells. Subsequently, we assessed the ability of LineageVAE to estimate time-series dynamics by comparing the estimated and observed cell states at each time point in the latent space. The data inferred by LineageVAE showed a similar spread to the observed data at the corresponding time points in the latent space (Fig. 2a, c and Supplementary Fig. 3). This indicates that LineageVAE effectively captured the time series transition of the cell states. We measured the distance between the inferred latent and observed cell states to quantitatively evaluate the accuracy of the inferred cell state transitions, at each time point using the silhouette score (Supplementary Fig. 2). Additionally, we employed an index that compares the inferred and observed latent cell states at each time point by counting the proportion of cells observed in the k neighborhood (here, k = 30). The silhouette score indicates that, As going back in time using this method, the silhouette score tends to be higher, and each cell tends to exhibit cohesion, resembling a common progenitor cell state. Furthermore, the ratio of the day 2 observed cells around an inferred cell increases on as one went back to the progenitor (Table. 1). This demonstrates that the model correctly traced time back to the undifferentiated region and captured changes in the cell state at each time point with high accuracy.

**Table 1:** The ratio of Observed time point in the k neighborhood. Observed time point of cells belonging to observed cell series in the latent space exists in the k neighborhood of each inferred cell series. The observation column is a control.

| Day | LineageVAE | Waddington-OT | Observation |
| --- | --- | --- | --- |
| day 2 | 2.20% | 22.10% | 3.70% |
| day 4 | 37.40% | 21.40% | 39.70% |
| day 6 | 60.30% | 56.60% | 56.60% |

### 3.3 Quantitative evaluation of accuracy and comparison with other methods for dynamics estimation

Subsequently, we compared the proposed method with other methods. First, we compared our method to pseudotime and scVelo (Trapnell *et al*., 2014) (Bergen *et al*., 2020, Fig. 2d and e). These two methods failed to predict the reagion of progenitor cells and the direction of differentiation. Moreover, they are widely used in differentiation research, and have contributed to the development of this field. However, when these methods are applied without time information for data from multiple observation times or for data measured simply after differentiation, sometimes accurately inferring the progenitor cell state may not be possible. The advantage of our method is the effective utilization of time information such as lineage tracing and observed time points, when valid information is obtained. A quantitative evaluation was performed by comparing the distances in the latent space between the mean point of day 2 observations and the progenitor cells predicted by each method (Table. 2). The LineageVAE performed better than pseudotime and scVelo in predicting differentiation trajectories and progenitor cell states.

**Table 2:** Distance from observed undifferentiated area. Each depict the Euclidean distance from the centroid of the observation on day 2 in the latent space. In the LineageVAE column, the distance represents the proximity to the progenitor cell inferred by LineageVAE. In the Pseudotime and scVelo columns, the distances correspond to the proximity to the rootcell of the trajectory inference in each respective method. These values serve as indices, indicating how far the cell state of the inferred progenitor is from the undifferentiated region.

| LineageVAE | Pseudotime | scVelo |
| --- | --- | --- |
| 1.46 | 3.84 | 3.66 |

Furthermore, we compared LineageVAE with another model that utilizes data with time information. We evaluated the performance of the model in the task of recovering a held-out time point. Specifically, we assessed the capacity of the model to accurately capture the marginal cell population on day 4 when trained solely on data from days 2 and 6. This evaluation utilized cells for which lineage tracing data were available. Second, we compared our method with that of Waddington-OT (Schiebinger *et al*., 2019), a technique that incorporates time point information (Fig. 2f-h). The data in Waddington-OT are embedded in the PCA space, and in LineageVAE are embedded in the latent space. Theredore, comparisons were difficult to make using the same distance scales. We employed an interpolation approach involving day 4 to gauge the performance. We projected the observed cells onto the same space and calculated the ratios using the adjacent observed time points of cells within the k-nearest neighbors (this time, k = 30) of the inferred day 4 cells. This approach aimed to mitigate the embedding effect caused by different dimensionality reductions inherent in the outcomes of each method. The simulated populations generated by our model outperformed those generated by the conventional method Waddington-OT.

### 3.4 LineageVAE transcriptome reconstruction accurately predicts RNA expression at unobserved time points

A trained LineageVAE model has the capability to generate transcriptomic measurements at each time point, tracing back to the progenitor state, based on post-differentiation transcriptomic observations. In this study, LineageVAE was used to estimate the transition of cell states over time in the latent space. Using these decoders, we reconstructed the transcriptome at each time point, effectively capturing the state transitions. The RNA expressed in each cell at each time point was sampled using the Poisson distributions, which allowed us to generate expression profiles for each lineage.

We analyzed marker gene expression by applying LineageVAE to the hematopoietic dataset to validate the biological accuracy of the reconstructed transcriptomes. We focused on lineages in which the cells displayed a single direction of differentiation and evaluated the expression of known markers associated with hematopoietic differentiation and markers for undifferentiated cells. We selected these marker genes using PanglaoDB, including post-differentiation markers (e.g., Sell and Itgam for monocytes) and undifferentiated markers (e.g., Cd34 for hematopoietic stem cells; Franzén *et al*., 2019). This analysis aimed to confirm whether the reconstructed transcriptomes aligned with the established biological knowledge. This analysis revealed a consistent pattern between the reconstructed transcriptomes and the expected biological behavior of the cells. Specifically, we observed a decreases in the expression of post-differentiation markers (Fig. 3a) and an increases in the expression of undifferentiated marker genes (Fig. 3b) as tracing back in time. In other cell types, gene expression at each time point was restored for many genes to align with the experimental facts (Supplementary Figs. 4, 5). These results further support the validity and accuracy of the reconstructed transcriptomes in capturing the dynamics of cell differentiation and the expected biological behavior of the cells. These findings suggest that LineageVAE can be used to infer transcriptome transitions.

**Figure 3:**
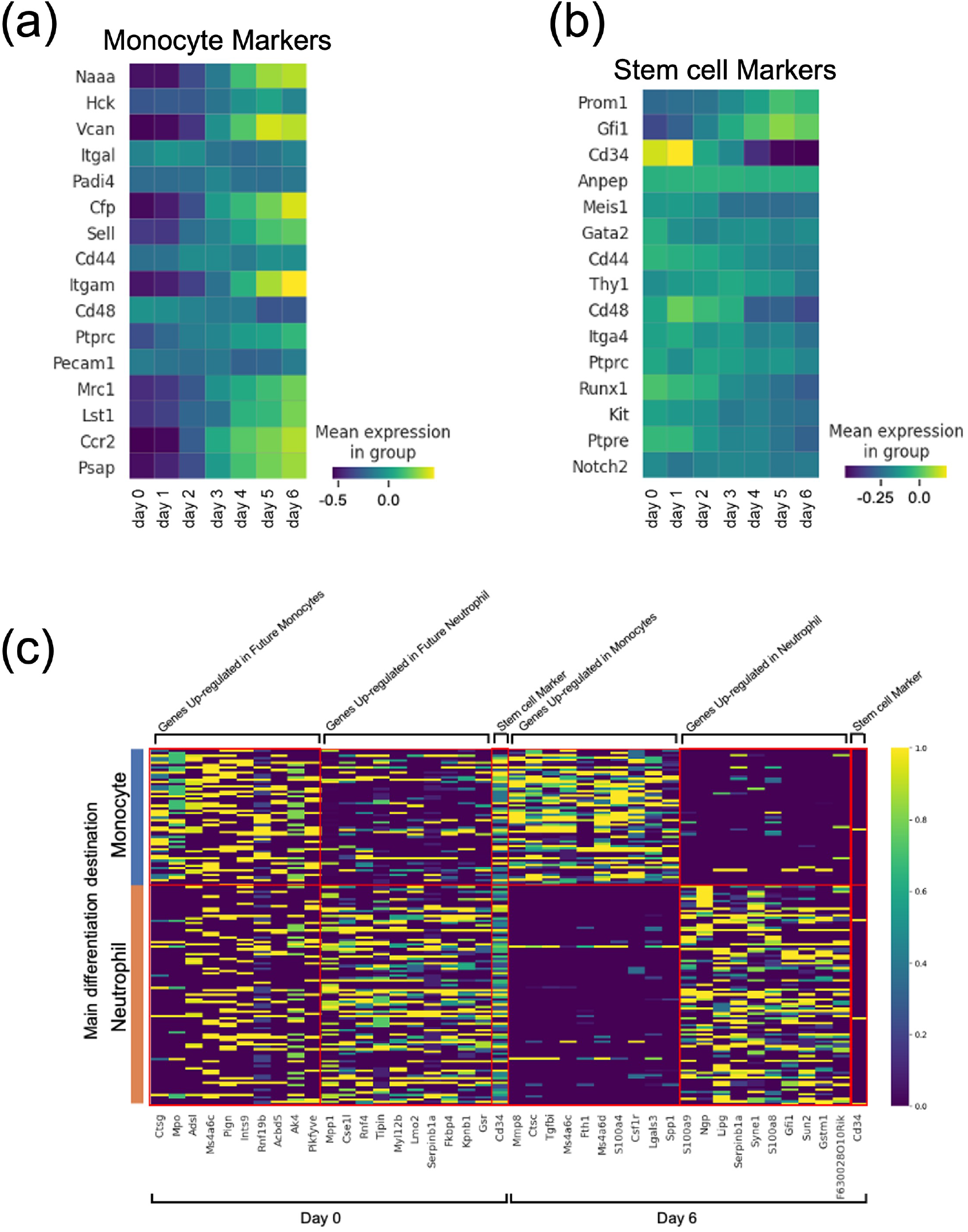
Reconstruction of historical expression by LineageVAE. **a**, Inferred expression of monocyte differentiation markers at each time point in cells that differentiate into monocytes. **b**, Inferred expression of undifferentiated marker at each time point in cells that differentiate into monocytes. **c**, Evaluation of progenitor bias by heatmaps of DEGs expression. Left: day 0, right: day 6.

### 3.5 Evaluation of unobservable ancestry bias through gene expression reconstruction

We subsequently discussed progenitor bias by taking advantage of the ability of the LineageVAE to reconstruct the transcriptome at each time point. First, we extracted the lineage that mainly differentiated into monocytes and neutrophils (lineage where 50% or more differentiated into a single cell type), and reconstructed the expression of each lineage progenitor (at day 0) from observation data at day 6 as model input data. Genes upregulated in the progenitor cells of the monocyte and neutrophil cell populations at each time point were selected by calculating differential expression genes (DEGs) of two populations, comparing their dispersion to the mean and standard deviation of dispersions within a specific bin of mean gene expression (Satija *et al*., 2015; Zheng et al., 2015) (Fig. 3c). The selected genes contained SerpinB1. SerpinB1 is reported as a critical protein in neutrophils, inhibiting serine proteases, such as NE, CG, and PR-3, to maintain mature neutrophil reserves in the bone marrow. Deficiency in SerpinB1 increases apoptosis and necrosis in the bone marrow, reducing neutrophil survival and resulting in fewer mobilizable neutrophils.

This study confirmed its high expression during neutrophil development, particularly at the promyelocyte stage. These findings support the accuracy of our model for predicting progenitor cell states (Benarafa *et al*., 2011). In addition to genes involved in blood cell differentiation, such as SerpinB1, many genes that are not known as marker genes were also included (Franzén *et al*., 2019). Some of these genes may reflect differentiation bias at an early stage. LineageVAE, which restores the expression of experimentally impossible-to-observe progenitor cells, may lead to discovering factors regulating differentiation at an early stage and may be useful for extracting indicator genes. Differential expression are calculated between monocytes and neutrophils in cells on day 6 and selected DEGs included the neutrophil marker Ngp. This also suggests that the candidates identified in these analyzes include genes that cause differentiation bias (Franzén *et al*., 2019).

### 3.6 Temporal differential changes in TF activity along differentiation trajectories

Our methodology enabled us to dissect the trajectory of differentiation trajectory and expression changes over time at the single cellular level. Here, we scrutinized the temporal variations in TF activity based on the reconstructed time series expression along this trajectory. TFs are proteins that control gene expression by binding to specific DNA sequences. They can either activate or repress the recruitment of RNA polymerases to genes (Fig. 4a). Initially, we summarized this regulatory relationship in a regulation matrix using previously reported information (Zhang *et al*., 2021). We assumed that the expression of TFs at a certain time point, denoted as t, controls the expression of target genes (TGs) at the next time point, t+1. We employed a generalized linear model (GLM) with TFs as the explanatory variables and TGs as the response variable (Fig. 4b). The weights, denoted by *w*_*tij*_, represent the learnable parameters in the GLM. Notably, these weights are intricately linked to the regulatory relationships between TF and TGs. When there is no regulatory relationship between a TFs and its TGs in this matrix, denoted by *r*_*tij*_ being 0, the corresponding weight *w*_*tij*_ is specifically set to 0 They are of each TF in regulating the expression of TGs for each gene is determined through regression. We defined the sum of these weight norms for each TF as their activity. First, to verify the TFs acitivity, using the trained model, we computed the average dynamics norm for each differentiation destination (Fig. 4c). This value is expected to increase significantly during the process of vigorous differentiation into mature cells. Notably, This suggests that the differentiation of monocytes and neutrophils is activated at an early time point, whereas the differentiation rate of basophils accelerates after a delayed time point. Overall, the total TF activity was increased before the norm of the dynamics increased (Supplementary Table 3). The activity of the TF Creb1 is assumed to increase from the initial stage in the cell group that differentiates into neutrophils (Fig. 4c). In the cell population that differentiates into neutrophils, the activity of the TF Creb1 is presumed to increase from an early stage before an increase in the norm of dynamics (Chen *et al*., 2023). Creb1 is a well-known TF that is crucial in regulating various aspects of neutrophil functions. This includes processes such as neutrophil extracellular traps (NETs) formation, phagocytic function, and the overexpression of pro-inflammatory cytokines and chemokines. In basophils, where differentiation acceleration was delayed, the timing of the increase in TFs activity was also delayed (Fig. 4d). With the ability to restore time-series expression, it is possible to evaluate TFs activity at each time point.

**Figure 4:**
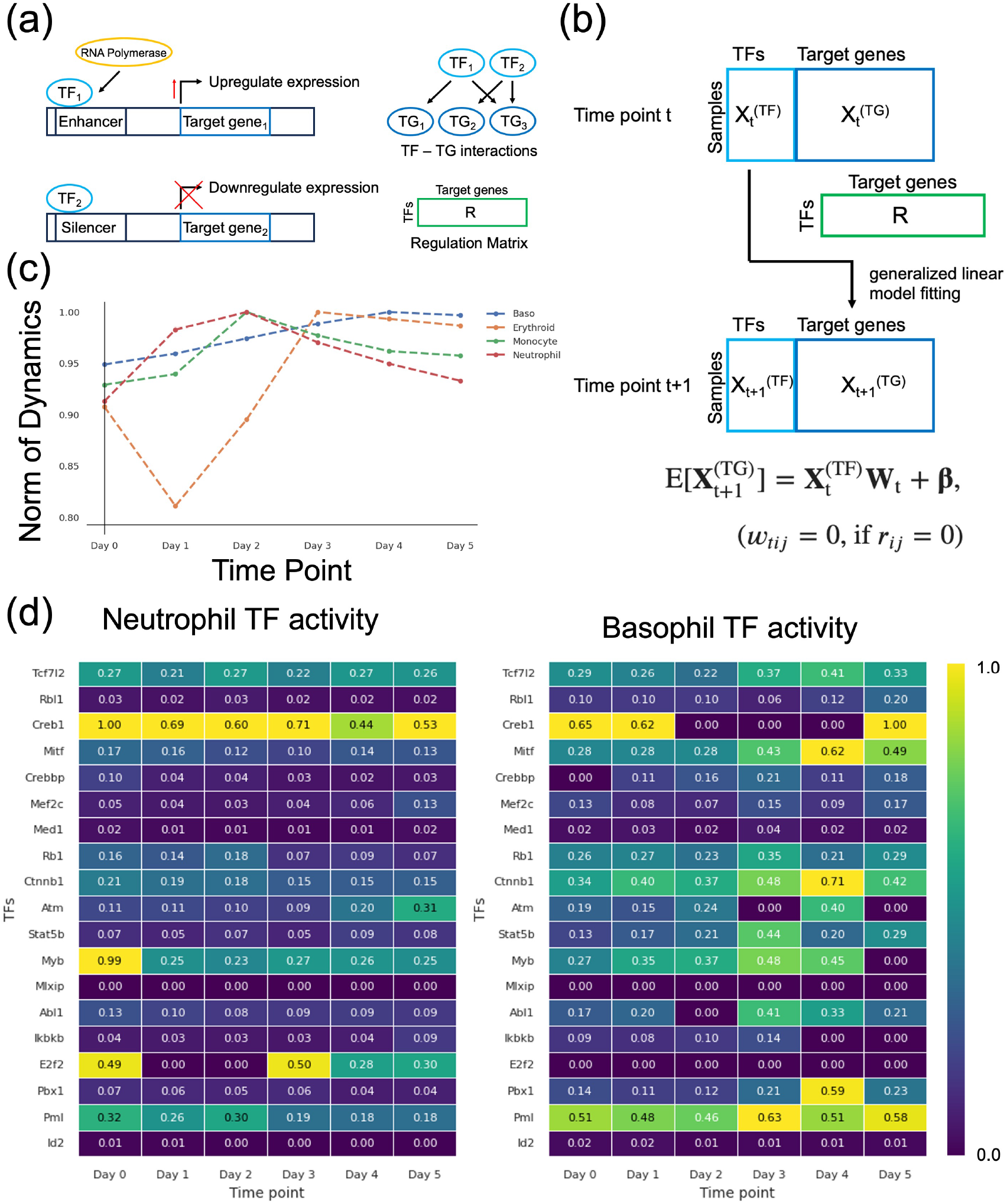
TF time differential activity analysis. **a**, Schematic diagram of how TF works. **b**, Application of GLM for TF activity analysis. TGs are downstream genes regulated by TFs. The relationship between TFs and TGs was calculated using the regulation matrix R.**c**, Norm of dynamics of each cell type. **d**, neutrophil (left) and basophil (right) time series TF activities. Among the genes used in the analysis, those reported in previous studies on the TFs-TGs regulatory relationship were selected as TFs.

### 3.7 LineagVI traces gene expression changes during the direct reprogramming from fibroblasts to induced endoderm progenitors at each time point

Finally, we applied LineageVAE to a lineage tracing dataset obtained through direct reprogramming using the CellTagging method (Biddy *et al*., 2018; Kamimoto *et al*., 2023), aiming to validate the robustness of the LineageVAE model. This study aimed to convert fibroblasts into induced endodermal progenitors (iEPs). We inputted the observed data from day 28 into the trained model, allowing us to estimate the state of each cell from days 0 to day 28. We subsequently compared these estimates with the actual observed data. The latent variables were projected onto the UMAP space, and the observation time, cell type, inferred time series cell state transitions of each cell, and time series cell state transitions of each cell in the two groups are displayed (Fig. 5a-d). The inferred time series data were retraced backwards in the latent space in a manner consistent with the actual observation data. This confirmed that the cells congregated in the fibroblast region as day 0 approached, suggesting that LineageVAE accurately predicted the past cell states. Cell type classification is based on quadratic programming, as detailed in the original study (Treutlein *et al*., 2016). Subsequently, we identified the lineages associated with iEP-enriched and iEP-depleted outcomes, similar to the approach used in the original study. Lineages achieving a 20–50% reprogramming success rate are designated as “iEP-enriched clones”. Conversely, lineages in which fewer than 3% of cells exhibit the characteristics of induced endodermal progenitors (iEPs) are categorized as “iEP-depleted clones”. We visualized the typical paths of iEP-enriched clones and iEP-depleted clones, excluding intermediate mixed lineages that existed within 0.1 times the standard deviation of the mean path of each group (Fig. 5d). One notable innovation of our method is its ability to reconstruct the expression intensities at each time point. The original study identified specific genes that were exclusively upregulated or downregulated following successful reprogramming or upon reaching a dead-end fate, as indicated by their high or low expression levels on day 28. We conducted additional analyses of the identified genes. By analyzing the historical gene expression changes for these specific genes, we compared the patterns between iEP-enriched and iEP-depleted clones (Fig. 5e). In this study, we applied time series k-means clustering to cluster time series expression changes and identified genes whose expression was enhanced in either the iEP-enriched or iEP-depleted groups (Tavenard *et al*., 2020). Consequently, we were able to infer the timing at which enhanced expression occurs during fate determination. Here, Apoa1 (an iEP marker) and Peg3 (which is crucial for the p53 apoptotic response) are presented as examples of genes that exhibit distinct patterns between the two groups (Fig. 5f and g). We estimated the gene expression levels for each lineage in an experimentally unobservable initial state (Fig. 5e, Supplementary Fig. 6). This indicates that Hes1, whose expression level has been reported to greatly affect growth and differentiation, expression level is significantly different between the iEP-enriched and iEP-depleted groups, suggesting the correctness of this inference (Castella *et al*., 2000; Yoshiura et al., 2007; Murata et al., 2005). Furthermore, gene regulatory network analysis can potentially enhance reprogramming yield by effectively targeting upstream transcription factors.

**Figure 5:**
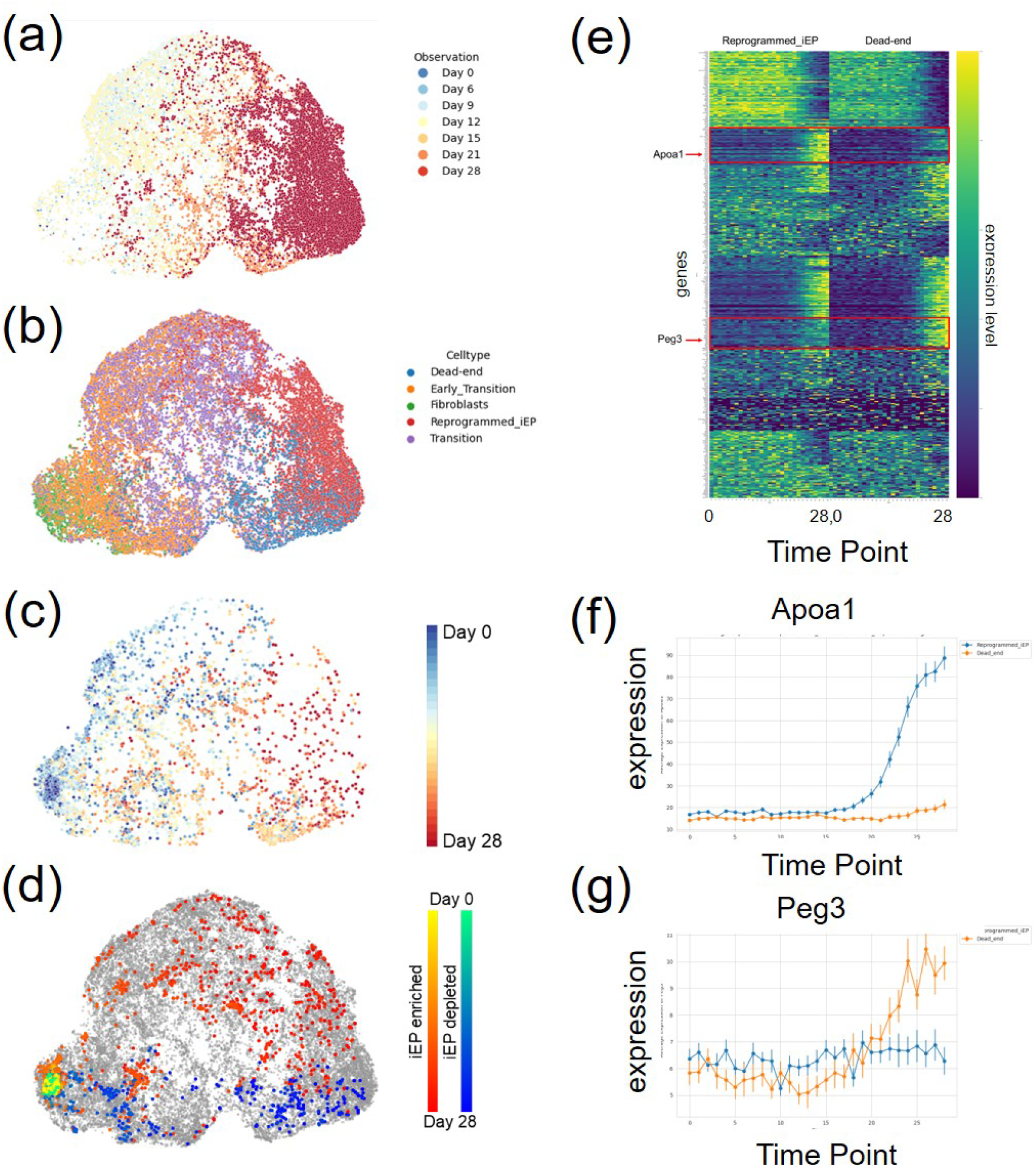
Application of LineagVAE to direct reprogramming dataset. **a-d**,UMAP visualization of the latent space inferred by the model. **a**, Time-series transition of cell state estimated by LineageVAE. **b**, Observed time points. **c**, Cell type annottations. **d**, Typical paths of iEP-enriched clones and iEP-depleted clones. **e**, Time-series changes of each gene expression in average of all cells. **f, g**, Restored expression along the time series. From top to bottom, Apoa1 (iEP marker), Peg3 (necessary for the p53 apoptotic response).

## 4 Discussion

We developed a VAE-based generative modeling framework for learning the dynamics in the latent space of time series scRNA-seq data. This model enables reasonable inference of unobserved cellular states and their lineage-specific stochastic dynamics in the latent space. Moreover, it allows for the recovery of the transcriptome data at each time point by inferring the technically unobservable progenitor state. Stem cell research, including induced pluripotent stem cells (iPS cells), actively seeks to advance regenerative medicine to restore lost functions by regenerating tissues and organs such as nerves, joints, muscles, skin, and the brain (Hoang *et al*., 2022; Yamanaka et al., 2012). Understanding the initial cellular state and the developmental mechanisms during the early differentiational stages is crucial for advancing regenerative therapies and organ transplantation by regulating differentiation into specific cell types. Most cancers are presumed to originate from a single cell with an oncogenic mutation, with additional mutations that facilitate tumor development(Greaves *et al*., 2012; Balani et al., 2017). However, the precise mechanisms underlying cancer initiation by a single transformed cell and the effects of additional mutations remain unclear(Vogelstein *et al*., 2013; Garraway *et al*., 2013). These analyses necessitate discussions that trace back to the progenitor cell state. However, lineage tracing has limitations, such as the ability to obtain observations only at discrete and sparse time points owing to scRNA-seq destroying cells during analysis, which makes it difficult to analyze the state of progenitor cells(Ding *et al*., 2013). Furthermore, lineage tracing experiments requires culturing to introduce the DNA barcode; scRNA-seq is performed after several rounds of cell division, and the direct progeny of the measured cells cannot be observed because of the destructive nature of the measurements. Regarding the aforementioned limitations, a variational autoencoder (VAE) is a powerful technique for capturing nonlinear latent structures in data (Kingma and Welling., 2013). Currently, there are VAE-based methods are available for single-cell data analysis, namely scVAE, scVI and scGEN (Grønbech., 2020; Romain., 2020; Lotfollahi., 2019). These methods have proven effective in capturing biological features within reduced dimensions. The diversity in their architectural designs renders VAEs well-suited for addressing several crucial challenges in scRNA-seq analysis, including dimensionality reduction, clustering, and data denoising (Wang *et al*., 2018; Geddes et al., 2019; Eraslan et al., 2019). Furthermore, models that combine the VAE and latent space vector arithmetics for high-dimensional single-cell gene expression data, captures the underlying structure of high-dimensional gene expression data in a lower-dimensional space, known as a manifold. The simple and linear nature of this latent space allows for linear extrapolations, which helps predict how gene expression changes in response to perturbations or other conditions. By utilizing the different vectors under different conditions, VAEs can effectively map these predicted changes back to the high-dimensional gene expression space, enabling the analysis and prediction of gene expression changes under various perturbations or conditions. Several methods have been developed using this property of the VAE characteristics, such as vicdyf and scMM (Minoura *et al*., 2019; Kojima et al., 2022). We gained insights from previous studies that applied the VAE to single-cell data analysis and hypothesized that by tracing back in time in the latent space, we could leverage the properties of the manifold space to recover a wide range of historical information. By estimating the dynamics in latent space, we inferred unobserved intermediate cellular states and common progenitors of all lineages. This method allows for the discussion of transcriptome and cell state transitions in dynamic processes. Furthermore, when observational data is limited, the VAE-based model remains valuable because it can generate pseudo cells from the latent space. This feature facilitates the analysis of dynamic biological processes. In addition, we accurately captured the phenomenon of branching into multiple cell types by effectively utilizing the property that cells sharing barcodes have identical progenitors and inferring back in time. From this, it seems reasonable to assume that multiple cells converge to a single progenitor state. By removing the regularization term that causes cells sharing a barcode to transition to a single-cell state, it may be possible to observe a reaction where the progenitors exhibit heterogeneity and diversity, transitioning into a homogeneous cell population post-reaction. For example, when a drug is added, multiple cells may transition into the same cell type. For downstream analysis, by restoring expression along the time series using LineageVAE, we considered gene regulation networks along the time axis. By considering the relationship between TF and TG, we were able to discuss regulatory lines in which variations in expression levels are involved in determining the direction of differentiation. When one wants to promote differentiation into a specific cell population, this will help guide the differentiation through the regulatory factors. We confirmed the robustness of the model by applying LineageVAE to a direct reprogramming dataset. Barcoding technology has developed rapidly in recent years, making it possible to create more detailed phylogenetic trees. Currently, our model considers only the absolute flow of time, however, by creating a model based on these phylogenetic relationships, it will be able to capture the unique flow of time for each differentiated area. Finally, the contribution of our proposal is that it can be optimized by assuming a single progenitor. Our model can be calculated only in the backward direction of time; however, creating a model that can be discussed both forward and backward in time may be possible by devising an optimization function and network. Until now, tracking time-series transitions in a single cell experimentally has been challenging owing to several limitations – each cell can be observed only once, and measurements can only be taken after the cell population has grown following barcode introduction. Our computational methodology addresses these experimental constraints, offering a solution to challenges that cannot be resolved using conventional methods in natural biotechnology. This model enables many analyses within the time series domain of single-cell studies, particularly in time series scenarios where data acquisition presents significant challenges.

## 5 Limitations of Study

LineageVAE is limited by its inability to predict future cell states. Additionally, accurately predicting the expression states beyond the observed region poses a challenge. These issues can be mitigated by advancing observation technologies through barcoding methods and utilizing mathematical models capable of conducting both forward and backward calculations. There is also room for consideration regarding the number of steps taken when tracing back time in the model (Bakken *et al*., 2020).

## 6 Method details

### 6.1 Generative model for sequential latent state transition

This section describes a generative model for time-series single-cell transcriptomes with lineage-tracing. This probabilistic model uses latent variables, *z*_*t*_ ∈ *R*^*m*^, where *m* is the dimension of the latent cell state space, *t* is the time of the cell state. Each lineage (a population of cells integrated with the same DNA barcode) begins with a unique progenitor cell state, which diffuses within the latent space over time. Expression was observed from the latent state after the cell state repeated diffusion until the observation time. The latent cell state is represented by ***z*** and follows a Gaussian prior distribution. Dynamics ***z***_*t*_− ***z***_*t*− 1_ represents the transition of the cell state during a certain period from time point ***t*** is expressed as the difference in the cell state between specific time points. These values followed a Gaussian prior distribution.

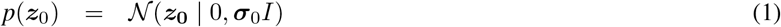

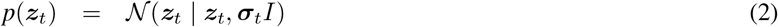

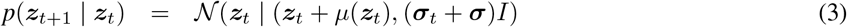

Where, ***σ***_0_ is optimizable parameter which defines the scale of the standard deviation of progenitor cell state, ***σ***_*t*_ represents the scale of the standard deviation of cell state at time point *t* and ***σ*** is optimizable parameter which defines the scale of the standard deviation in dynamics. We note that this distribution can be chosen arbitrarily and can include options such as a Gaussian distribution with mean ***z***_initial observation_ parameterized by the observed latent state of early undifferentiated cell populations and optimizable parameter *σ*, for example, as follows to facilitate optimization.

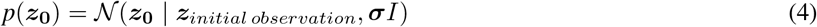

We modeled the latent cell state transition after a certain period as dynamics ***z***_*t*_− ***z***_*t*−1_. This generative model assumes that the time evolution of the latent cell state ***z*** follows the Wiener process. For each lineage, By assuming an initial latent cell state ***z***_**0**_, we constructed a generative model for each transcriptome ***X*** at time point *T*_0_ as follows.

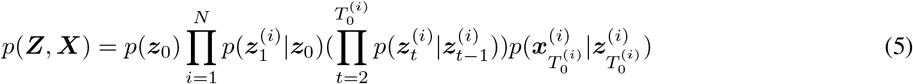

Where 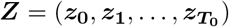.

### 6.2 Variational inference of transcriptome dynamics for time-series lineage-tracing single-cell data

We developed a computational methodology for estimating the dynamics of a time series lineage tracing single-cell dataset. Suppose we have a dataset *X* consists of spliced and unspliced transcriptomes of a single cell ***s***∈*R*^*g*^ and ***u*** ∈ *R*^*g*^, respectively, where *g* is the number of genes, lineage information represented by a barcode tag, and time point information representing when the data were observed. As a variational inference of the aforementioned generative model, we encode the cell state and infer the transition of the cell state going back in time. Our methodology stochastically embeds the single cell transcriptome of each cell into a latent cell state space using a deep neural network. Our methodology assumes a variational posterior distribution of the cell state at the previous time point for each latent cell state, and then recursively predicts progenitor cells. In the following sections, we describe the generative model for transcriptomes and the variational approximation of the posterior distributions of latent variables with lineage information in the following sections.

### 6.3 Variational autoencoder for embedding high-dimensional transcriptome into low-dimensional latent space

To embed transcriptome data into a low-dimensional latent space, similar to previous studies, we used the variational autoencoder (VAE) framework (Way *et al*., 2018; Lopez et al., 2018; Lotfollahi., 2019; Grønbech., 2020; Nagaharu *et al*., 2022). A VAE is a deep generative neural network that reduces the dimensionality and generates data. Let ***x*** be the data and ***z*** be the set of low-dimensional latent variables, *z* ∈ *R*^*m*^ where *m* is the dimension of the latent cell state space, the VAE consists of an encoder and decoder neural network that parameterizes the variational posterior *q*_Φ_(***z***| ***x***) and likelihood *p*_Θ_(***x***|***z***), respectively. VAE replaces the true intractable posterior *p*(***z***|***x***) with a variational posterior *q*_Φ_(***z***|***x***) and approximates the intractable integrals with *p*_Θ_(***x***|***z***), which is a likelihood of the data given a sample from the variational posterior. This approach allows the encoder to estimate the low-dimensional latent variables ***z*** from the data *p*(***x***) and the decoder to learn the generation of data from a given low-dimensional representation. The VAE objective function is the lower bound of the marginal likelihood of ***x*** (evidence lower bound; ELBO), which can be written with a reconstruction term and a Kullback-Leibeler (KL) divergence regularization term:

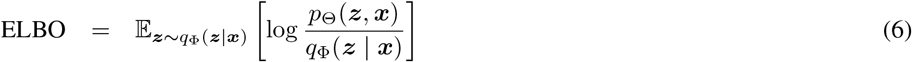

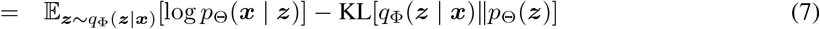

This probabilistic model assumes that the prior over the latent variables *p*(***z***) is typically chosen as an isotropic standard multivariate Gaussian distribution.

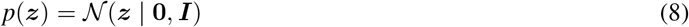

We assumed that the transcriptome ***x*** follows the Poisson distribution.

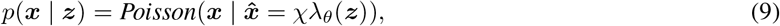

where χ∈ *R* is the mean expression across all genes in the single cell and λ_*θ*_(***z***) ∈ *R*^*g*^ is the decoding neural network of the latent cell state with 50 hidden units, two layers, and layer normalization.

### 6.4 Variational inference of the latent variable posterior

To estimate the time series latent cell state ***z*** and corresponding transition dynamics over a specific time period, we assumed that the variational distribution of dynamics ***z***_***t***_− ***z***_***t*−1**_ is a Gaussian distribution dependent on the latent cell state ***z***_***t***_ as shown below:

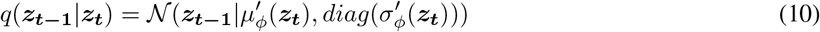

where *µ*^*′*^ and *σ*^*′*^ are neural networks with 50 hidden units, two layers, and layer normalization. These formulations correspond to the assumption that the reverse time evolution of the latent cell state ***z*** follows an discrete advectiondiffusion model.

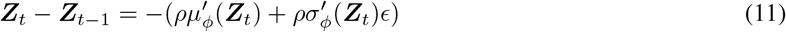

where 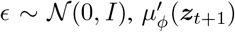 is correspond to the average of latent state, dynamics 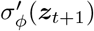 is correspond to the fluctuation of latent state dynamics and *ρ* is a parameter that indicates that the absolute value of the dynamics, which is the transition from the previous time, is sufficiently smaller than the variation of the cell state in the latent space at that time, and it can be optimizable. Additionally, we performed variational inference backward in time in the latent space, starting from the point corresponding to the observation time of the progenitor cell state.

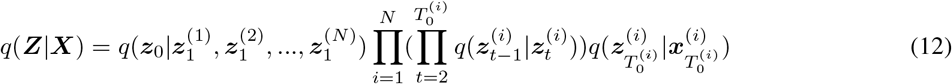

The key idea of LineageVAE is to assume that cells with the same lineage are in a uniquely identical state ***z***_**0**_ at time 0, and that cell states change from ***z***_**1**_ into ***z***_**0**_ follows the same dynamics as the variational distribution at other time points in the latent space so that ***z***_**0**_ is represented as follows:

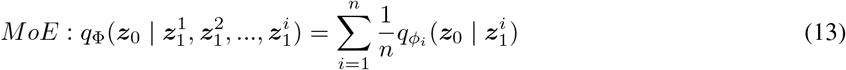

For the estimation process of the parameters of the generative models and the variational distribution, we maximized the *ELBO* as defined below:

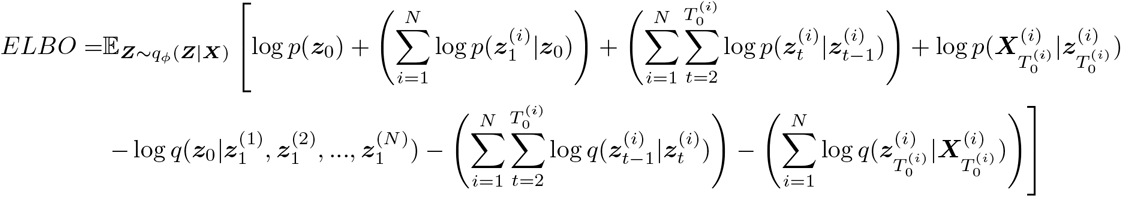

The calculation of *ELBO* necessitates sampling from the variational distribution. When sampling ***z***_0_, we sampled 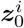 from 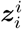 and and selectively chose one of them, given that LineageVAE assumes a single progenitor cell states from each lineage. Optionally, to optimize efficiency, ***z***_0_ can be chosen using stratified sampling. During optimization, the latent space series ***z*** is derived through reparameterized sampling from ***z*** inferred in the variational processes. Where *E*_*p*(*x*)_[*f* (*x*)] represents the expectation of *f* (*x*) given *x*∼*P* (*x*). For this maximization, we used the Adam optimizer with a learning rate of 0.0001, mini batch size of 30 composed of a single lineage, and 1000 epochs at total. In the first step, we estimated encoder parameter *ϕ* for embedding into low-dimensional latent space. In the second step, we fixed *ϕ* and estimated encoder parameter *ϕ*^*′*^ for encoding dynamics and decoder parameter *θ*. All implementations were based on the PyTorch library of Python language.

### 6.5 Optimization using micro and macro information constraints

Our study employed lineage tracing information to infer macroscopic cell state transitions. Additionally, by differentiating between spliced and unspliced mRNAs in standard scRNA-seq data, following the same approach as existing methodologies (La Manno *et al*., 2018; Nagaharu et al., 2022), it is possible to infer the cell state change during a micro-duration around the observed cell state. Here, we assumed that the mean parameters of spliced and unspliced transcriptomes followed the differential equation of splicing kinetics as with existing tools for RNA velocity estimation:

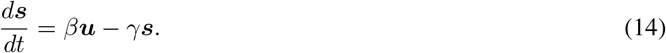

where *β*∈*R*^*g*^ is gene-specific splicing rates of unspliced transcriptome and *γ*∈*R*^*g*^ is a vector of gene-specific degradation rates of spliced transcriptome. The changes in the spliced transcriptome were expressed as:

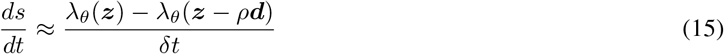

From equations (14) and (15), we derived the mean parameter of the unspliced transcriptome as follows:

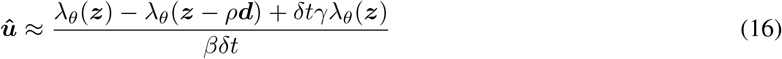

We assumed that the transcriptome ***u*** has the following Poisson distribution approximately.

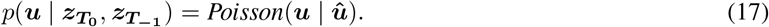

Considering the micro constraints at the observed time point, *ELBO* can be expressed as follows:

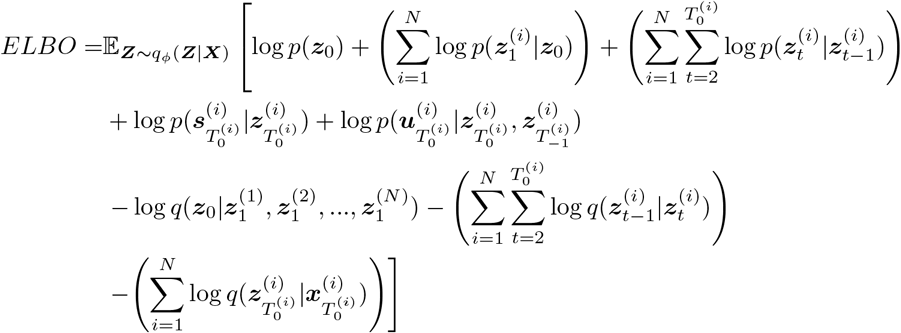

### 6.6 Model architecture and optimization

Optimization was conducted using the Adam optimizer with AMSGrad (Reddi *et al*., 2018). The neural network structure comprised two hidden layers with 50 hidden units each. The learning rate was set as 0.0001. For the LARRY data, mini batch sizes of 30 were employed, with each mini batch loaded with data corresponding to a single barcode. Learning was performed out over 200 epochs.

### 6.7 Data preprocessing

We first selected the 1000 most variable genes for the transcriptome count data by applying the scanpy *highly_variable_genes* function in log-normalized counts. Raw counts were used as model inputs.

### 6.8 Visualization of latent representations

The mean parameters for the variational posteriors were used as the latent variables. Latent variables obtained from trained models were visualized on the two-dimensional space using the “umap” package in Python.

### 6.9 Evaluation of biological meaning of embedding using Euclidean distance

We calculated the average coordinates and their standard deviations in the latent space for each cell type, which were manually annotated based on the gene expression in the original paper. We then calculated the Euclidean distance and p-value between the cell types and confirmed that the same cell types were embedded in close positions in the latent space.

### 6.10 Evaluation of embedding accuracy using observation data

To evaluate the accuracy of the time series estimation in a latent space containing data of various degrees of differentiation, we counted the observed dates of cells in the k-neighborhood of each inferred cell in the observation space. We calculated the ratio of observed dates of observed cells existing in the k-neighborhood. We used it as an index of the similarity to the observed cells at a certain time point and degree of differentiation.

### 6.11 Comparison of embedding accuracy using observation data

To compare the accuracy of the methods, we interpolated day 4 using the data from days 2 and 6. Notably, WaddingtonOT method operates within the PCA space, whereas LineageVAE operates within the latent space. This distinction makes it challenging to directly compare the two methods using the same distance scale. To evaluate the performance of these methods, we developed an interpolation strategy focused on day 4. This strategy involved calculating ratios using the observation data from observed time points (this time, days 2, 4, and 6) for cells within K (this time 30) neighborhood cells around each estimated day 4 cell in each embedded space. The primary objective of this approach was to address and mitigate the embedding differences inherent in the outcomes generated by each of the two methods.

### 6.12 TFs activity evaluation

Here, we assumed that the expression of TFs (*i*) at a certain time point, denoted as t, controls the expression of TGs (*j*) at the next time point, t+1. We employed a GLM with TFs as explanatory variables and TGs as the response variable. Let 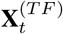 be the matrix representing the TFs expression of each sample at time point *t*, 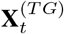 be the matrix representing the TGs expression of each sample at time point *t*, and **R** be a matrix summarizing the TF-TG interactions. Each element of the matrix is denoted by *r*_*ij*_. Let weights *W* be the learnable parameters determined by regression. Each element of the matrix is denoted by *w*_*tij*_. The following regression model was used in this study.

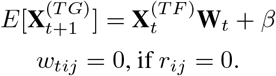

To evaluate the TF activity, we calculated the sum of the absolute values of these weights. This value was then scaled between 0 and 1 to eliminate the influence of the total amount of expression detected. A absolute value of each element in the set *w*_*ti*_, where *j* represents the various elements in the set. The TF activity was then computed as the sum of these absolute values. To normalize the TF activity within a standardized range of 0 to 1, each TF activity value was divided by the maximum TF activity value calculated across all TF *i* instances. In conclusion, we defined the TF Activity for TF *i* at time point *t* as:

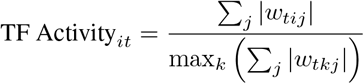

## 8 Acknowledgements

This research was funded by multiple sources. The Grants-in-Aid for Scientific Research (B) (grant no. 20H04281), Grants-in-Aid for Scientific Research on Innovative Areas on Information Physics of Living Matters (grant no. 22H04839), Grant-in-Aid for Transformative Research Areas (platforms for Advanced Technologies and Research Resources) (grant no. 22H04925), Grant-in-Aid for Transformative Research Areas (A) (grant no. 23H04938), and Grant-in-Aid for Research Activity Start-up (grant no. 20K22839) were provided by the Japan Society for the Promotion of Science (JSPS). Additional support was received from RADDAR-J (grant no. JP22ek0109488), the Project for P-PROMOTE (grant nos. JP22ama221215 and JP22ama221501), Brain/MINDS Health and Diseases (grant no. JP22wm0425007), the Interdisciplinary Cutting-edge Research (grant no. JP23wm0325068), and the Advanced Genome Research and Bioinformatics Study to Facilitate Medical Innovation (GRIFIN) (grant no. JP23tm0424226) from the Japan Agency for Medical Research and Development (AMED). The Moonshot Moonshot R&D program (grant no. JPMJMS2025) and ACT-X program (grant no. JPMJAX20AB) also contributed, through the Japan Science and Technology Agency (JST). Further support came from the Medical Research Center Initiative for High Depth Omics and Multilayered Stress Diseases at Tokyo Medical and Dental University. Supercomputing resources were provided by the Shirokane supercomputer at the Human Genome Center of the University of Tokyo, the TSUBAME3.0 supercomputer at the Tokyo Institute of Technology, and the AI Bridging Cloud Infrastructure (ABCI) at the National Institute of Advanced Industrial Science and Technology (AIST).

## 9 Author contributions

K. Minoura conceived the idea for this study. K. Majima designed and performed the experiments under the supervision of Y.K. and T.S. All authors have read and approved the final manuscript.

## 11 Resource availability

### 11.1 Lead contact

Further information and resource requests should be directed to, and will be fulfilled by, the lead contact, Teppei Shimamura

### 11.2 Materials availability

This study did not generate new unique reagents.

### 11.3 Data and code availability

The data from experiments of Weinreb et al. were downloaded from https://github.com/AllonKleinLab/paper-data/blob/master/Lineage_tracing_on_transcriptional_landscapes_links_state_to_fate_during_differentiation/README.md (commit: d8f0969). Data for the Biddy et al. were downloaded from GEO (GSE99915). The LineageVAE model was implemented in Python using the PyTorch deep learning library, and the code is available at https://github.com/LzrRacer/LineageVAE/. All the original codes have been deposited at Zenodo and are publicly available on publication. The DOIs are listed in the Key Resources Table.

Any additional information required to reanalyze the data reported in this paper is available from the lead contact upon request.

**Table S1:** Batch Size and Cell Types Included.

| Batch Size | 10 | 20 | 30 | 40 | 50 |
| --- | --- | --- | --- | --- | --- |
| Undifferentiated | 573 | 443 | 324 | 232 | 158 |
| Neutrophil | 490 | 383 | 262 | 184 | 120 |
| Monocyte | 382 | 323 | 257 | 208 | 126 |
| Mast | 53 | 2 | 2 | 2 | 1 |
| Baso | 49 | 24 | 2 | 2 | 1 |
| Meg | 12 | 4 | 1 | 0 | 0 |
| Erythroid | 10 | 7 | 0 | 0 | 0 |
| Eos | 5 | 2 | 1 | 1 | 1 |
| Lymphoid | 2 | 1 | 1 | 0 | 0 |
| Ccr7_DC | 1 | 1 | 0 | 0 | 0 |
| Number of selected cells | 1410 | 1180 | 870 | 720 | 500 |

**Table S2:**
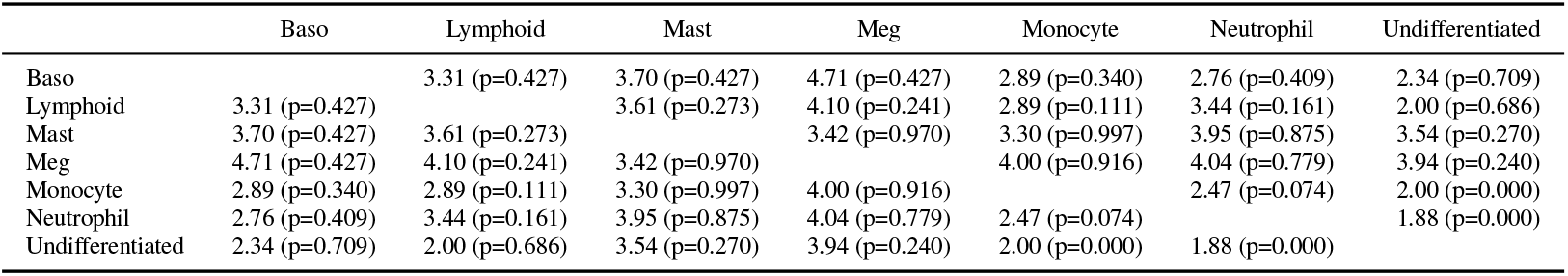
Average distance and p-value for each cell type. The mean and standard deviation on the latent space were calculated for each cell type. This table displays the average distance and p-value for each cell type.

**Table S3:** TF Activity per time point for Different Cell Types. The calculated activities of the defined TFs for each day were summed, and the totals for each date were then normalized, ensuring that the day with the highest sum equaled 1.00.

| Cell Type | Day |  |  |  |  |  |
| --- | --- | --- | --- | --- | --- | --- |
|  | 0 | 1 | 2 | 3 | 4 | 5 |
| Baso | 0.76 | 0.80 | 0.63 | 0.94 | 1.00 | 0.86 |
| Monocyte | 1.00 | 0.79 | 0.67 | 0.68 | 0.80 | 0.98 |
| Neutrophil | 1.00 | 0.67 | 0.69 | 0.74 | 0.72 | 0.81 |

**Figure S1:**
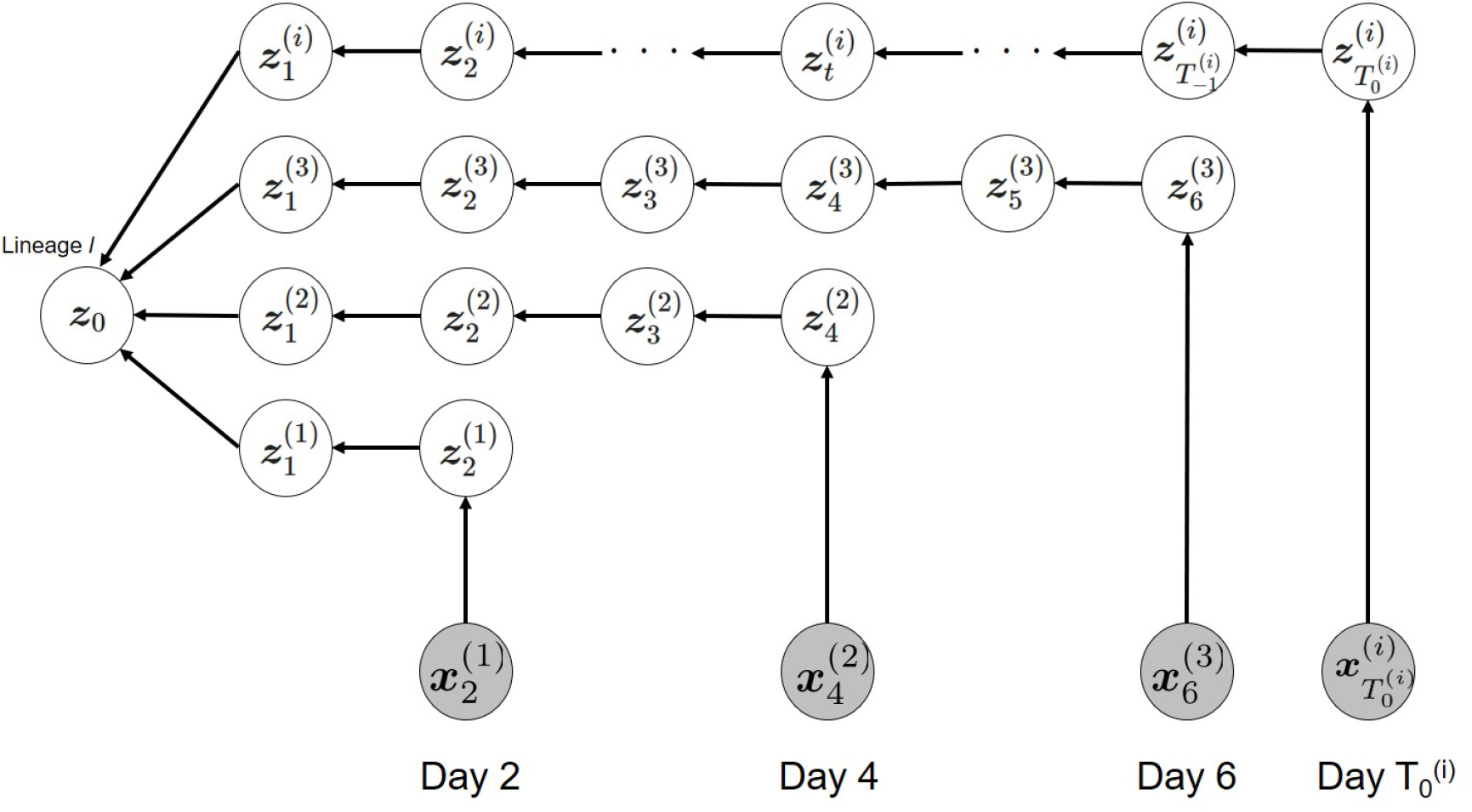
LineageVAE graphical model. Shaded nodes 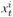 indicate observed data, and white node 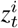 symbolizes latent variables. Edges indicate dependencies. Variational inference is conducted backward in time from the observed data.

**Figure S2:**
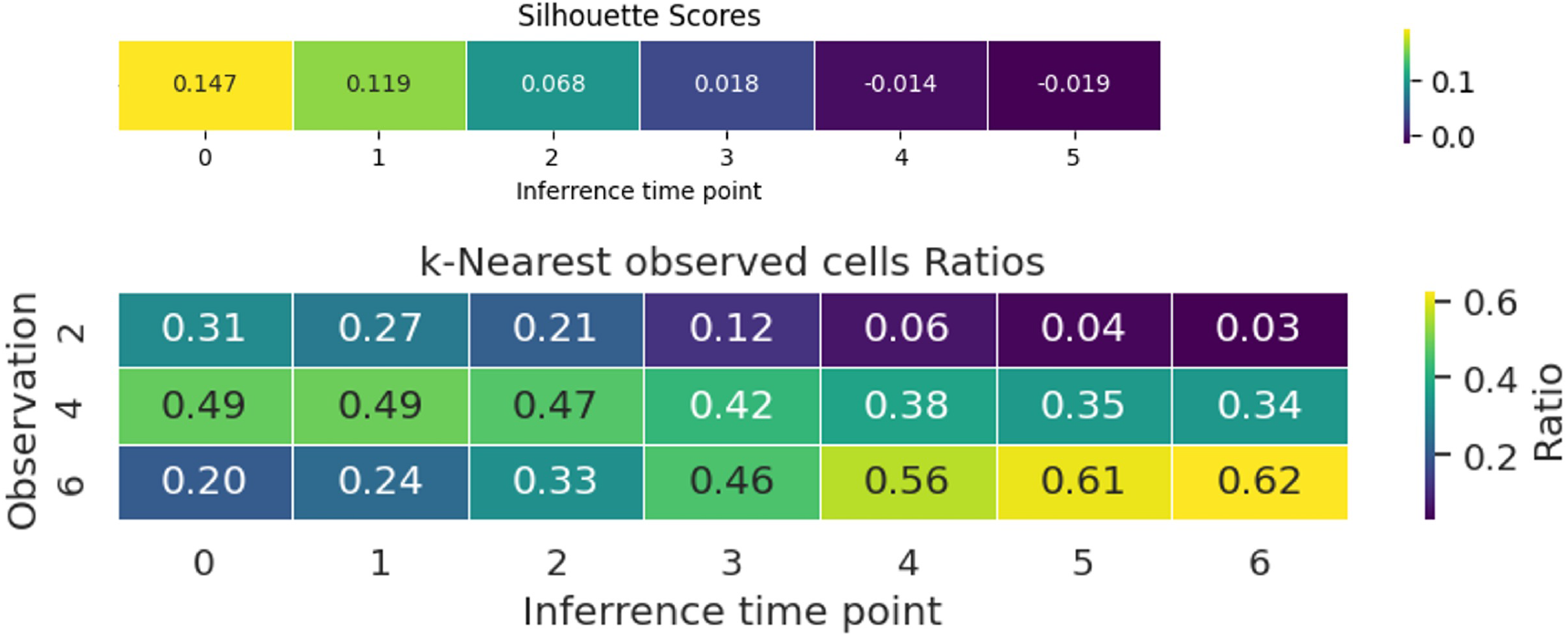
Quantitative evaluation of time series inferrence. **a** Silhouette score for cells in each estimated time point for observed cells. **b** Ratio of observed cells that are k neighbors of the inferred cells.

**Figure S3:**
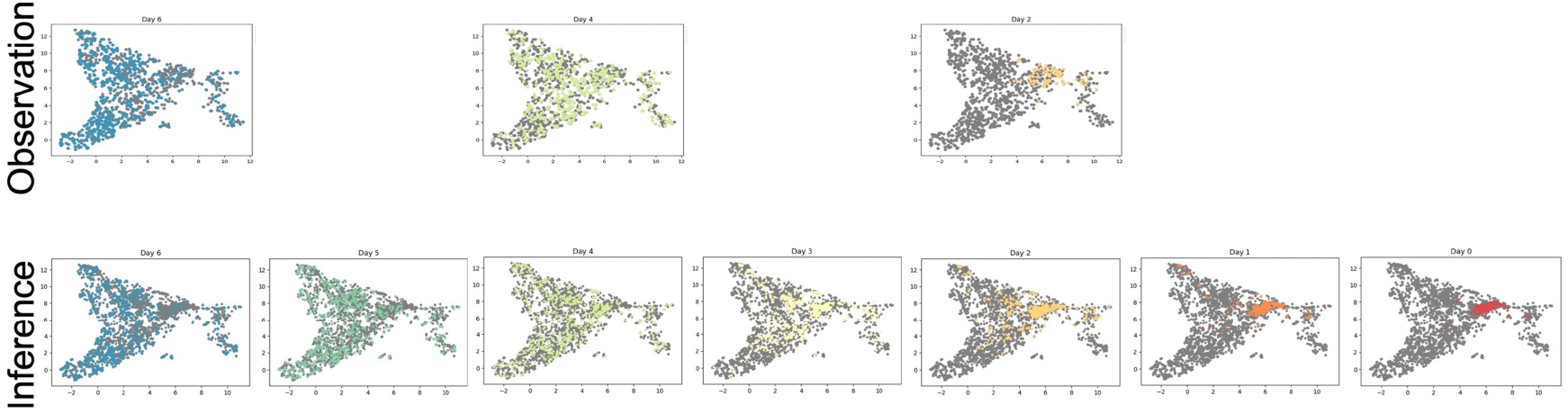
Overview of the time series inferrence. (Upper) Observation cells and observation time points (Lower) Inferred cells and time points

**Figure S4:**
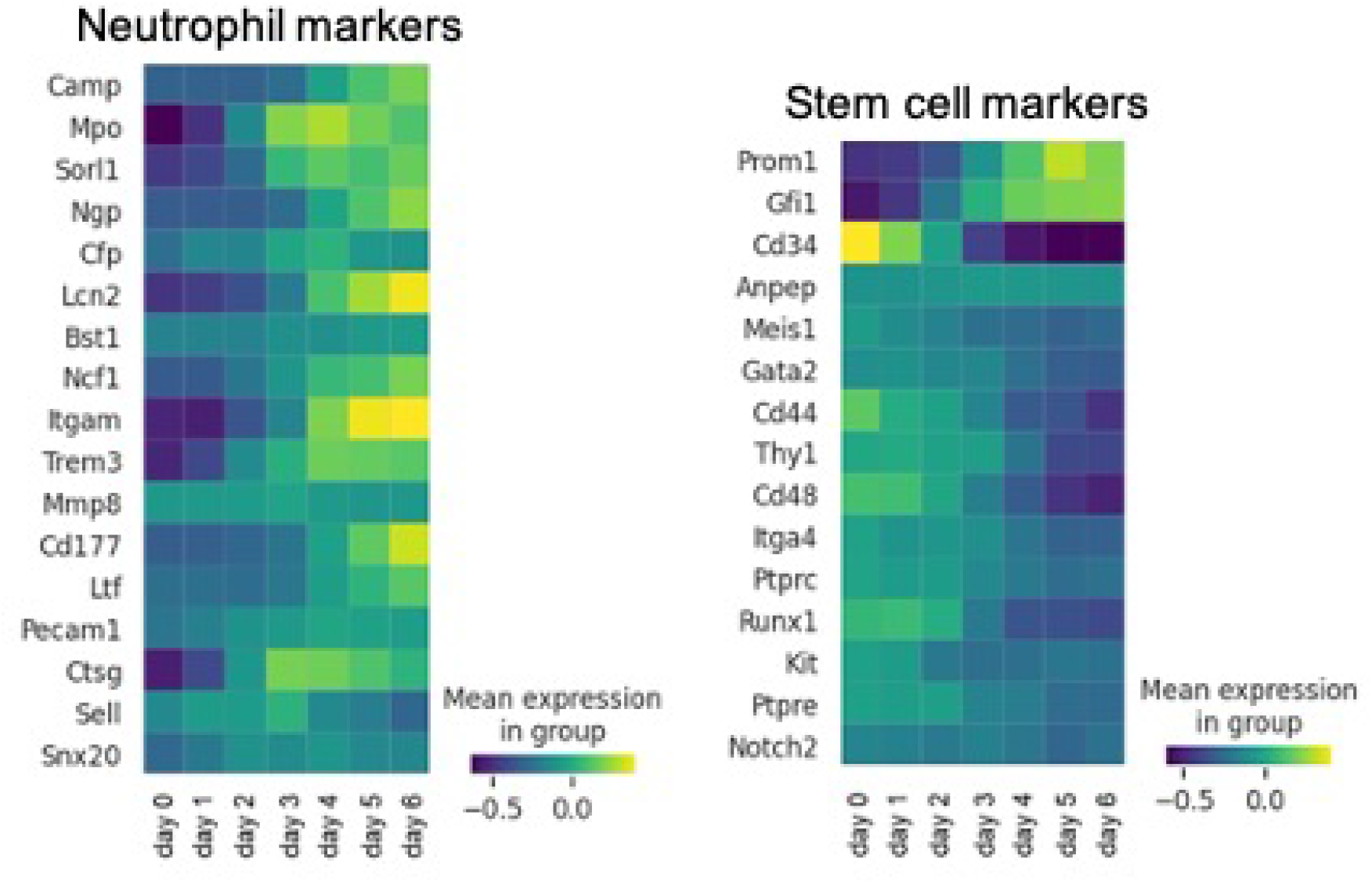
Reconstruction of historical expression by LineageVAE. (left) Inferred expression of Neutrophil differentiation markers at each time point in cells that differentiate into Neutrophil. (right) Inferred expression of undifferentiated marker at each time point in cells that differentiate into Neutrophil.

**Figure S5:**
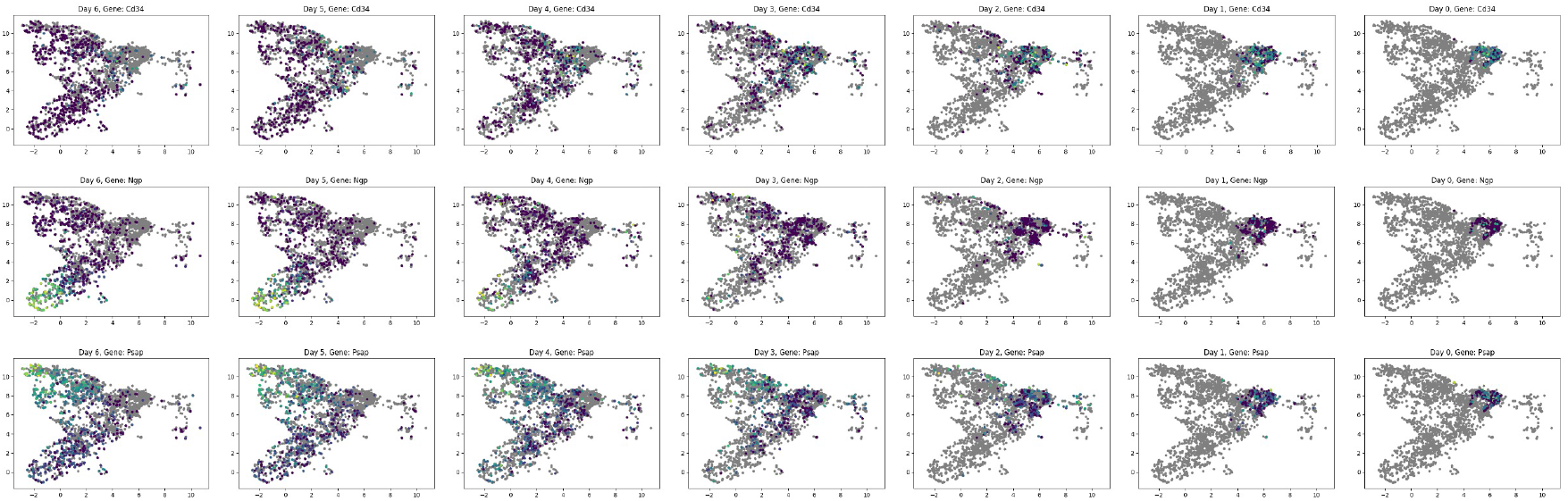
Visualization of recovered gene expression on UMAP. (Upper) Neutrophil Marker: Ngp expression at each time point. (Middle) Monocyte Marker: Psap expression at each time point. (Lower) undifferentiated marker: Cd34 expression at each time point.

**Figure S6:**
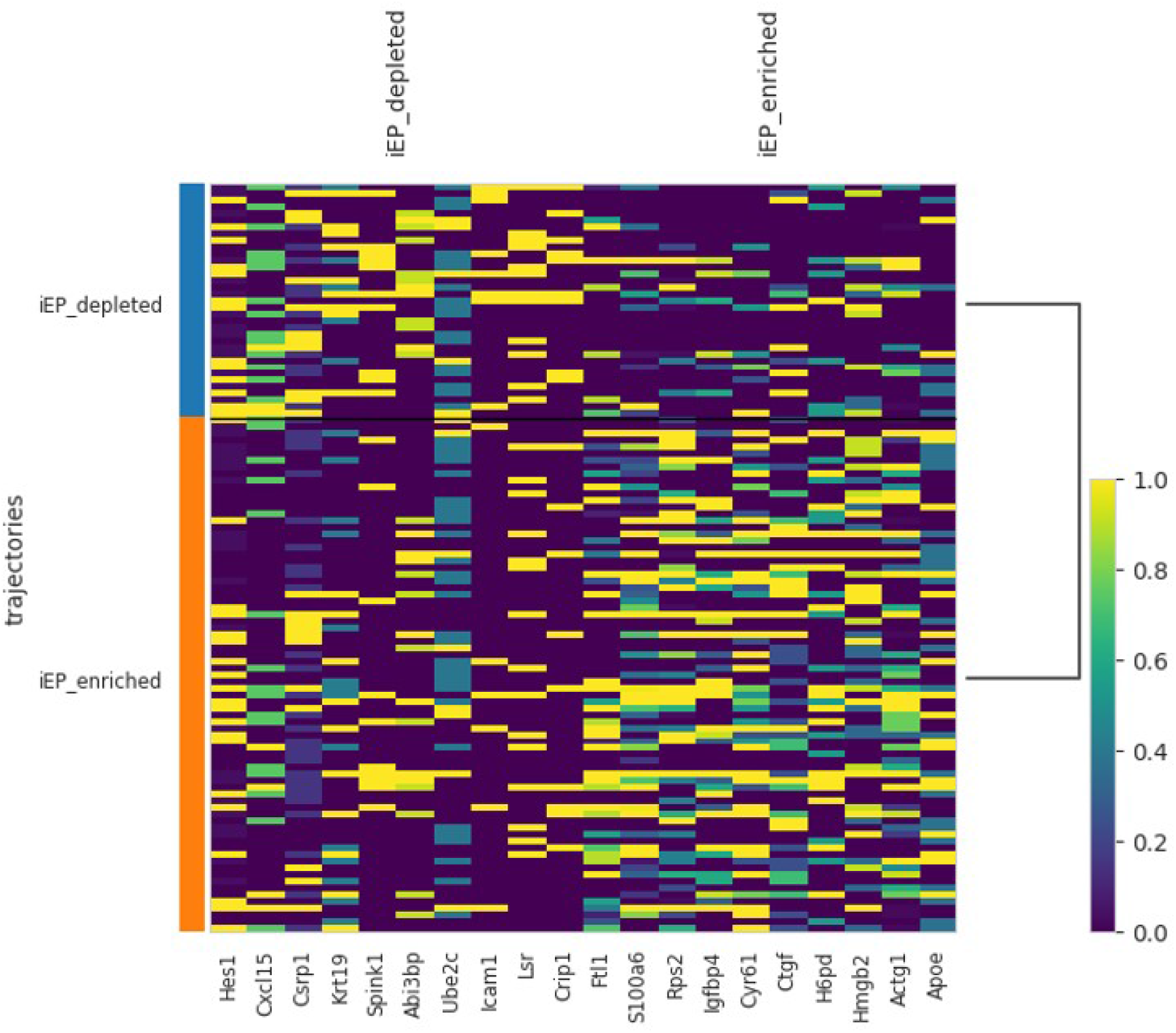
Evaluation of progenitor bias by heatmaps. DEGs expression on day 0, which is experimentally unobservable after reprogramming induction.

## Reference

Hoang DM, Pham PT, Bach TQ, Ngo ATL, Nguyen QT, Phan TTK, Nguyen GH, L. PTT, Hoang VT, Forsyth NR, Heke M, Nguyen LT. Stem cell-based therapy for human diseases. Signal Transduct Target Ther. 2022 Aug 6;7(1):272.

Yamanaka S. Induced pluripotent stem cells: past, present, and future. Cell Stem Cell. 2012 Jun 14;10(6):678–684.

Greaves M, Maley CC. Clonal evolution in cancer. Nature. 2012 Jan 18;481(7381):306–13. doi: 10.1038/nature10762.

Balani S, Nguyen LV, Eaves CJ. Modeling the process of human tumorigenesis. Nat Commun. 2017 May 25;8:15422. doi: 10.1038/ncomms15422.

Vogelstein B, Papadopoulos N, Velculescu VE, Zhou S, Diaz LA Jr, Kinzler KW. Cancer genome landscapes. Science. 2013 Mar 29;339(6127):1546–58.

Garraway LA, Lander ES. Lessons from the cancer genome. Cell. 2013 Mar 28;153(1):17–37.

Ding J, Sharon N, Bar-Joseph Z. Temporal modelling using single-cell transcriptomics. Nat Rev Genet. 2022 Jun;23(6):355–368.

Hwang, B., Lee, J.H., Bang, D. (2018). Single-cell RNA sequencing technologies and bioinformatics pipelines. Experimental and Molecular Medicine, 50, 1–14.

Trapnell C, Cacchiarelli D, Grimsby J, Pokharel P, Li S, Morse M, Lennon NJ, Livak KJ, Mikkelsen TS, Rinn JL. The dynamics and regulators of cell fate decisions are revealed by pseudotemporal ordering of single cells. Nat Biotechnol. 2014; 32(4):381.

Saelens W, Cannoodt R, Todorov H, Saeys Y. A comparison of single-cell trajectory inference methods. Nat Biotechnol. 2019; 37(5):547–54.

Song, D., Li, J.J. PseudotimeDE: inference of differential gene expression along cell pseudotime with well-calibrated p-values from single-cell RNA sequencing data. 2021; Genome Biol 22, 124.

La Manno, G., Soldatov, R., Zeisel, A., Braun, E., Hochgerner, H., Petukhov, V., Lidschreiber, K., Kastriti, M. E., Lönnerberg, P., Furlan, A., Fan, J., Borm, L. E., Liu, Z., van Bruggen, D., Guo, J., He, X., Barker, R., Sundström, E., Castelo-Branco, G., Cramer, P., Adameyko, I., Linnarsson, S., Kharchenko, P. V. (2018). RNA velocity of single cells. Nature, 560, 494–498.

Bergen, V., Lange, M., Peidli, S., Wolf, F. A., Theis, F. J. (2020). Generalizing RNA velocity to transient cell states through dynamical modeling. Nat Biotechnol 38, 1408–1414.

Hashimoto, T., Gifford, D., and Jaakkola, T. (2016). Learning population-level diffusions with generative RNNs. In Proceedings of the 33rd International Conference on Machine Learning, B. Maria Florina and Q.W. Kilian, eds. (PMLR).

Fischer, D.S., Fiedler, A.K., Kernfeld, E.M., Genga, R.M.J., Bastidas-Ponce, A., Bakhti, M., Lickert, H., Hasenauer, J., Maehr, R., and Theis, F.J. (2019). Inferring population dynamics from single-cell RNA-sequencing time series data. Nat. Biotechnol. 37, 461–468.

Weinreb, C., Rodriguez-Fraticelli, A., Camargo, F.D., Klein, A.M. (2020). Lineage tracing on transcriptional landscapes links state to fate during differentiation. Science. 367, eaaw3381

Oren, Y., Tsabar, M., Cuoco, M.S., Amir-Zilberstein, L., Cabanos, H.F., Hütter, J.C., Hu, B., Thakore, P.I., Tabaka, M., Fulco, C.P., Colgan, W., Cuevas, B.M., Hurvitz, S.A., Slamon, D.J., Deik, A., Pierce, K.A., Clish, C., Hata, A.N., Zaganjor, E., Lahav, G., Politi, K., Brugge, J.S., Regev, A.(2021). Cycling cancer persister cells arise from lineages with distinct programs. Nature. 596, 576–582.

Wang SW, Herriges MJ, Hurley K, Kotton DN, Klein AM. CoSpar identifies early cell fate biases from single-cell transcriptomic and lineage information. Nat Biotechnol. 2022 Jul;40(7):1066–1074.

Forrow A, Schiebinger G. LineageOT is a unified framework for lineage tracing and trajectory inference. Nat Commun. 2021 Aug 16;12(1):4940.

Wang K, Hou L, Wang X, Zhai X, Lu Z, Zi Z, Zhai W, He X, Curtis C, Zhou D, Hu Z. PhyloVelo enhances transcriptomic velocity field mapping using monotonically expressed genes. Nat Biotechnol. 2023 Jul 31.

Kingma, D.P., and Welling, M. (2013). Auto-Encoding Variational Bayes. Arxiv.

Grønbech, C.H., Vording, M.F., Timshel, P.N., Sønderby, C.K., Pers, T.H., Winther, O. (2020). scVAE: variational auto-encoders for single-cell gene expression data. Bioinformatics. 36, 4415–4422

Romain, L., Regier, J., Cole, M.B., Jordan, M.I., Yosef, N. (2018). Deep generative modeling for single-cell transcriptomics. Bioinformatics. 36, 4415–4422

Lotfollahi M, Wolf FA, Theis FJ. scGen predicts single-cell perturbation responses. Nat Methods. 2019 Aug;16(8):715–721.

Akiba, T., Sano, S., Yanase, T., Ohta, T. and Koyama, M. (2019). Optuna: A Next-generation Hyperparameter Optimization Framework. Arxiv.

Bakken, T.E., Jorstad, N.L., Hu, Q., Lake, B.B., Tian, W., Kalmbach, B.E., Crow, M., Hodge, R.D., Krienen, F.M., Sorensen, S.A., et al. (2020). Evolution of cellular diversity in primary motor cortex of human, marmoset monkey, and mouse. Biorxiv 2020.03.31.016972.

Cao, J., Cusanovich, D.A., Ramani, V., Aghamirzaie, D., Pliner, H.A., Hill, A.J., Daza, R.M., McFaline-Figueroa, J.L., Packer, J.S., Christiansen, L., et al. (2018). Joint profiling of chromatin accessibility and gene expression in thousands of single cells. Science 361, eaau0730.

Chen, S., Lake, B.B., and Zhang, K. (2019). High-throughput sequencing of the transcriptome and chromatin accessibility in the same cell. Nat Biotechnol 37, 1452–1457.

Gayoso, A., Steier, Z., Lopez, R., Regier, J., Nazor, K.L., Streets, A., and Yosef, N. (2021). Joint probabilistic modeling of single-cell multi-omic data with totalVI. Nat Methods 1–11.

González-Blas, C.B., Minnoye, L., Papasokrati, D., Aibar, S., Hulselmans, G., Christiaens, V., Davie, K., Wouters, J., and Aerts, S. (2019). cisTopic: cis-regulatory topic modeling on single-cell ATAC-seq data. Nat Methods 1–4.

Hao, Y., Hao, S., Andersen-Nissen, E., Mauck, W.M., Zheng, S., Butler, A., Lee, M.J., Wilk, A.J., Darby, C., Zager, M., et al. (2021). Integrated analysis of multimodal single-cell data.

Haghverdi, L., Lun, A.T.L., Morgan, M.D., and Marioni, J.C. (2018). Batch effects in single-cell RNA-sequencing data are corrected by matching mutual nearest neighbors. Nat Biotechnol 36, 421.

Hu, Y., Huang, K., An, Q., Du, G., Hu, G., Xue, J., Zhu, X., Wang, C.-Y., Xue, Z., and Fan, G. (2016). Simultaneous profiling of transcriptome and DNA methylome from a single cell. Genome Biol 17, 88.

Jin, S., Zhang, L., and Nie, Q. (2020). scAI: an unsupervised approach for the integrative analysis of parallel single-cell transcriptomic and epigenomic profiles. Genome Biol 21, 25.

Joost, S., Annusver, K., Jacob, T., Sun, X., Dalessandri, T., Sivan, U., Sequeira, I., Sandberg, R., and Kasper, M. (2020). The Molecular Anatomy of Mouse Skin during Hair Growth and Rest. Cell Stem Cell 26, 441–457.e7.Cell.

Kennedy, M.K., Willis, C.R., and Armitage, R.J. (2006). Deciphering CD30 ligand biology and its role in humoral immunity. Immunology 118, 143–152.

Levine, J.H., Simonds, E.F., Bendall, S.C., Davis, K.L., Amir, E.D., Tadmor, M.D., Litvin, O., Fienberg, H.G., Jager, A., Zunder, E.R., et al. (2015). Data-Driven Phenotypic Dissection of AML Reveals Progenitor-like Cells that Correlate with Prognosis. Cell 162, 184–197.

Lopez, R., Regier, J., Cole, M.B., Jordan, M.I., and Yosef, N. (2018). Deep generative modeling for single-cell transcriptomics. Nat Methods 15, 1053–1058.

Ma, S., Zhang, B., LaFave, L.M., Earl, A.S., Chiang, Z., Hu, Y., Ding, J., Brack, A., Kartha, V.K., Tay, T., et al. (2020). Chromatin Potential Identified by Shared Single-Cell Profiling of RNA and Chromatin. Cell.

McInnes, L., Healy, J., and Melville, J. (2018). UMAP: Uniform Manifold Approximation and Projection for Dimension Reduction. Arxiv.

Reddi, S.J., Kale, S., and Kumar, S. (2019). On the Convergence of Adam and Beyond. Arxiv.

Robert, C. P. and Casella, G. (2004). Monte Carlo Statistical Methods. Springer Texts Statistics 511–543 doi:10.1007/978-1-4757-4145-213.

Rodriques, S.G., Stickels, R.R., Goeva, A., Martin, C.A., Murray, E., Vanderburg, C.R., Welch, J., Chen, L.M., Chen, F., and Macosko, E.Z. (2019). Slide-seq: A scalable technology for measuring genome-wide expression at high spatial resolution. Science 363, 1463–1467.

Rosa, F.D., and Pabst, R. (2005). The bone marrow: a nest for migratory memory T cells. Trends Immunol 26, 360–366.

Schep, A.N., Wu, B., Buenrostro, J.D., and Greenleaf, W.J. (2017). chromVAR: inferring transcription-factor-associated accessibility from single-cell epigenomic data. Nat Methods 14, 975–978.

Shi, Y., Siddharth, N., Paige, B., and Torr, P.H.S. (2019). Variational Mixture-of-Experts Autoencoders for Multi-Modal Deep Generative Models. Arxiv.

Schiebinger G, Shu J, Tabaka M, Cleary B, Subramanian V, Solomon A, Gould J, Liu S, Lin S, Berube P, Lee L, Chen J, Brumbaugh J, Rigollet P, Hochedlinger K, Jaenisch R, Regev A, Lander ES. Optimal-Transport Analysis of Single-Cell Gene Expression Identifies Developmental Trajectories in Reprogramming. Cell. 2019 Feb 7;176(4):928–943.e22.

Sønderby, C. K., Raiko, T., Maaløe, L., Sønderby, S. K. and Winther, O. (2016). Ladder Variational Autoencoders. Arxiv.

Stuart, T., Butler, A., Hoffman, P., Hafemeister, C., Papalexi, E., Mauck, W.M., Hao, Y., Stoeckius, M., Smibert, P., and Satija, R. (2019). Comprehensive Integration of Single-Cell Data. Cell 177, 1888–1902.e21.

Stuart, T., Srivastava, A., Lareau, C., and Satija, R. (2020). Multimodal single-cell chromatin analysis with Signac. Biorxiv 2020.11.09.373613.

Sun, S., Zhu, J., Ma, Y. & Zhou, X. Sun, S., Zhu, J., Ma, Y., and Zhou, X. (2019). Accuracy, robustness and scalability of dimensionality reduction methods for single-cell RNA-seq analysis. Genome Biol 20, 269.

Svensson, V. (2020). Droplet scRNA-seq is not zero-inflated. Nat Biotechnol 38, 147–150.

Svensson, V., Gayoso, A., Yosef, N., and Pachter, L. (2020). Interpretable factor models of single-cell RNA-seq via variational autoencoders. Bioinformatics 36, 3418–3421.

Welch, J.D., Kozareva, V., Ferreira, A., Vanderburg, C., Martin, C., and Macosko, E.Z. (2019). Single-Cell Multi-omic Integration Compares and Contrasts Features of Brain Cell Identity. Cell 177, 1873-1887.e17.

Wu, M., and Goodman, N. (2018). Multimodal Generative Models for Scalable Weakly-Supervised Learning. Arxiv.

Xiong, L., Xu, K., Tian, K., Shao, Y., Tang, L., Gao, G., Zhang, M., Jiang, T., and Zhang, Q.C. (2019). SCALE method for single-cell ATAC-seq analysis via latent feature extraction. Nat Commun 10, 4576.

Zhu, C., Yu, M., Huang, H., Juric, I., Abnousi, A., Hu, R., Lucero, J., Behrens, M.M., Hu, M., and Ren, B. (2019). An ultra high-throughput method for single-cell joint analysis of open chromatin and transcriptome. Nat Struct Mol Biol 26, 1063–1070.

Zhu, C., Preissl, S., and Ren, B. (2020). Single-cell multimodal omics: the power of many. Nat Methods 17, 11–14.

Zuo, C., and Chen, L. (2020). Deep-joint-learning analysis model of single cell transcriptome and open chromatin accessibility data. Brief Bioinform bbaa287-.

Franzén O. Gan, LM. and Björkegren, JLM. (2019). PanglaoDB: a web server for exploration of mouse and human single-cell RNA sequencing data.

Biddy, B.A., Kong, W., Kamimoto, K., Guo, C., Waye, S.E., Sun, T, Morris, S.A., (2018) Single-cell mapping of lineage and identity in direct reprogramming. Nature 564, 219–224.

Kamimoto K, Adil MT, Jindal K, Hoffmann CM, Kong W, Yang X, Morris SA. Gene regulatory network reconfiguration in direct lineage reprogramming. Stem Cell Reports. 2023 Jan 10;18(1):97–112.

Tavenard, R., Faouzi, J., Vandewiele, G., Divo, F., Androz, G., Holtz, C., Payne, M., Yurchak, R., Rußwurm, M., Kolar, K., Woods, E. (2020). Tslearn, A Machine Learning Toolkit for Time Series Data. Journal of Machine Learning Research, 21(118), 1–6.

Castella P, Sawai S, Nakao K, Wagner JA, Caudy M. HES-1 repression of differentiation and proliferation in PC12 cells: role for the helix 3-helix 4 domain in transcription repression. Mol Cell Biol. 2000 Aug;20(16):6170–83.

Yoshiura S, Ohtsuka T, Takenaka Y, Nagahara H, Yoshikawa K, Kageyama R. Ultradian oscillations of Stat, Smad, and Hes1 expression in response to serum. Proc Natl Acad Sci U S A. 2007 Jul 3;104(27):11292–7.

Murata K, Hattori M, Hirai N, Shinozuka Y, Hirata H, Kageyama R, Sakai T, Minato N. Hes1 directly controls cell proliferation through the transcriptional repression of p27Kip1. Mol Cell Biol. 2005 May;25(10):4262–71.

Way GP, Greene CS. Extracting a biologically relevant latent space from cancer transcriptomes with variational autoencoders. Pac Symp Biocomput.2018;23:80–91.

Nagaharu K, Kojima Y, Hirose H, Minoura K, Hinohara K, Minami H, Kageyama Y, Sugimoto Y, Masuya M, Nii S, Seki M, Suzuki Y, Tawara I, Shimamura T, Katayama N, Nishikawa H, Ohishi K. A bifurcation concept for B-lymphoid/plasmacytoid dendritic cells with largely fluctuating transcriptome dynamics. Cell Rep. 2022 Aug 30;40(9):111260.

Wang D, Gu J. VASC: Dimension Reduction and Visualization of Single-cell RNA-seq Data by Deep Variational Autoencoder. Genomics Proteomics Bioinformatics. 2018 Oct;16(5):320–331.

Geddes TA, Kim T, Nan L, Burchfield JG, Yang JYH, Tao D, Yang P. Autoencoder-based cluster ensembles for single-cell RNA-seq data analysis. BMC Bioinformatics. 2019 Dec 24;20(Suppl 19):660.

Eraslan G, Simon LM, Mircea M, Mueller NS, Theis FJ. Single-cell RNA-seq denoising using a deep count autoencoder. Nat Commun. 2019 Jan 23;10(1):390. doi: 10.1038/s41467-018-07931-2.

Minoura K, Abe K, Nam H, Nishikawa H, Shimamura T. A mixture-of-experts deep generative model for integrated analysis of single-cell multiomics data. Cell Rep Methods. 2021 Sep 15;1(5):100071. doi: 10.1016/j.crmeth.2021.100071.

Satija R, Farrell JA, Gennert D, Schier AF, Regev A. Spatial reconstruction of single-cell gene expression data. Nat Biotechnol. 2015 May;33(5):495–502.

Zheng GX, Terry JM, Belgrader P, Ryvkin P, Bent ZW, Wilson R, Ziraldo SB, Wheeler TD, McDermott GP, Zhu J, Gregory MT, Shuga J, Montesclaros L, Underwood JG, Masquelier DA, Nishimura SY, Schnall-Levin M, Wyatt PW, Hindson CM, Bharadwaj R, Wong A, Ness KD, Beppu LW, Deeg HJ, McFarland C, Loeb KR, Valente WJ, Ericson NG, Stevens EA, Radich JP, Mikkelsen TS, Hindson BJ, Bielas JH. Massively parallel digital transcriptional profiling of single cells. Nat Commun. 2017 Jan 16;8:14049.

Benarafa C, LeCuyer TE, Baumann M, Stolley JM, Cremona TP, Remold-O’Donnell E. SerpinB1 protects the mature neutrophil reserve in the bone marrow. J Leukoc Biol. 2011 Jul;90(1):21–9. doi: 10.1189/jlb.0810461. Epub 2011 Jan 19.

Zhang Z, Deng Y, Zheng G, Jia X, Xiong Y, Luo K, Qiu Q, Qiu N, Yin J, Lu M, Liu H, Gu Y, He Z. SRGN-TGFβ2 regulatory loop confers invasion and metastasis in triple-negative breast cancer. Oncogenesis. 2017 Jul 10;6(7):e360.

Korpetinou A, Skandalis SS, Labropoulou VT, Smirlaki G, Noulas A, Karamanos NK, Theocharis AD. Serglycin: at the crossroad of inflammation and malignancy. Front Oncol. 2014 Jan 13;3:327.

Kolset SO, Pejler G. Serglycin: a structural and functional chameleon with wide impact on immune cells. J Immunol. 2011 Nov 15;187(10):4927–33.

Latchman DS. Transcription factors: an overview. Int J Exp Pathol. 1993 Oct;74(5):417–22.

Zhang Y, Liu T, Hu X, Wang M, Wang J, Zou B, Tan P, Cui T, Dou Y, Ning L, Huang Y, Rao S, Wang D, Zhao X. CellCall: integrating paired ligand-receptor and transcription factor activities for cell-cell communication. Nucleic Acids Res. 2021 Sep 7;49(15):8520–8534.

Chen J, Bai Y, Xue K, Li Z, Zhu Z, Li Q, Yu C, Li B, Shen S, Qiao P, Li C, Luo Y, Qiao H, Dang E, Yin W, Gudjonsson JE, Wang G, Shao S. CREB1-driven CXCR4hi neutrophils promote skin inflammation in mouse models and human patients. Nat Commun. 2023 Sep 22;14(1):5894.

Treutlein B, Lee QY, Camp JG, Mall M, Koh W, Shariati SA, Sim S, Neff NF, Skotheim JM, Wernig M, Quake SR. Dissecting direct reprogramming from fibroblast to neuron using single-cell RNA-seq. Nature. 2016 Jun 16;534(7607):391–5.

